# Phosphorylation barcode-dependent signal bias of the dopamine D1 receptor

**DOI:** 10.1101/2020.03.03.971846

**Authors:** Ali I. Kaya, Nicole A. Perry, Vsevolod V. Gurevich, T.M. Iverson

## Abstract

Agonist-activated G protein-coupled receptors (GPCRs) must correctly select from hundreds of potential downstream signaling cascades and effectors. To accomplish this, GPCRs first bind to an intermediary signaling protein, such as G protein or arrestin. These intermediaries initiate signaling cascades that promote the activity of different effectors, including several protein kinases. The relative roles of G proteins versus arrestins in initiating and directing signaling is hotly debated, and it remains unclear how the correct final signaling pathway is chosen given the ready availability of protein partners. Here, we begin to deconvolute the process of signal bias from the dopamine D1 receptor (D1R) by exploring factors that promote the activation of ERK1/2 or Src, the kinases that lead to cell growth and proliferation. We found that ERK1/2 activation involves both arrestin and Gαs, while Src activation depends solely on arrestin. Interestingly, we found that the phosphorylation pattern influences both arrestin and Gαs coupling, suggesting an additional way the cells regulate G protein signaling. The phosphorylation sites in the D1R intracellular loop 3 are particularly important for directing the binding of G protein versus arrestin and for selecting between the activation of ERK1/2 and Src. Collectively, these studies correlate functional outcomes with a physical basis for signaling bias and provide fundamental information on how GPCR signaling is directed.

**Significance Statement:** The functional importance of receptor phosphorylation in GPCR regulation has been demonstrated. Over the past decade, the phospho-barcode concept was developed to explain the multi-dimensional nature of the arrestin-dependent signaling network downstream of GPCRs. Here, we used the dopamine-1 receptor (D1R) to explore the effect of receptor phosphorylation on G protein-dependent and arrestin-dependent ERK and Src activation. Our studies suggest that D1R intracellular loop-3 phosphorylation affects both G proteins and arrestins. Differential D1R phosphorylation can direct signaling toward ERK or Src activation. This implies that phosphorylation induces different conformations of receptor and/or bound arrestin to initiate or select different cellular signaling pathways.

## Introduction

The classical paradigm of G protein-coupled receptor (GPCR) signaling posits that agonist-activated GPCRs bound G proteins, which in turn stimulated effectors that regulate second messengers and initiate a cellular response. To terminate G protein-dependent signaling, GPCR kinases (GRKs) phosphorylate active GPCRs at their carboxyl terminus and/or intracellular loops, thereby priming them for arrestin binding (1). Arrestins bind to phosphorylated and activated receptors with high affinity, which simultaneously prevents further G-protein coupling (desensitization), and initiates receptor internalization. Later it was discovered that receptor-associated arrestins also direct signaling toward alternative pathways. In arrestin-mediated signaling, which is almost exclusively supported by the non-visual arrestins (i.e., arrestin-2 and arrestin-3^1^), arrestins act as scaffolds or signal adapters (2-6), binding simultaneously to activated and phosphorylated GPCRs and downstream effectors.

Although it is tempting to divide GPCR signaling into these two branches, G protein-dependent and arrestin-dependent, the nature of signaling in the cell is much more complex (reviewed in (1, 7, 8)). As a result, this classical paradigm does not fully explain the range of biological responses observed when a GPCR is activated. For example, agonist identity can affect the recruitment of different G-proteins or arrestins, which determines the selection of downstream effectors (9). In addition, some activated GPCRs may interact directly with non-canonical signaling partners, including Src-family kinases (10-21). There are also numerous studies suggesting a specific role for the non-visual arrestins in receptor-independent activation of ERK1/2, AKT, JNK3, and NF-κB (6, 22-26). Complexity of the signaling process is further increased by cross-talk between the signaling pathways initiated by G proteins and arrestins (27). When one considers the sheer number of receptors, ligands, effectors, and alternative pathway connections, it is not surprising that the best efforts of the field have not yet been able to identify how an extracellular stimulus, be it hormonal or sensory, directs signaling toward the correct biological response.

The complexity of GPCR signaling is well exemplified by the five dopamine receptors (D1R – D5R). Dopamine receptors support the neural plasticity required for a myriad of physiological functions (28-30), particularly learning and memory (31). Hijacking of this reward pathway by drugs of abuse contributes to addiction (31). Studies of D1R have already identified Gs and G-olf as cognate G proteins that transduce D1R signaling (28, 32). In addition, the role of arrestin-3 in signaling from the dopamine receptors has been demonstrated using knockout mice (33-37) and cultured cells (38). This arrestin-3-mediated signaling contributes to the pathology of addiction, as suggested by studies showing that the effects of amphetamine and apomorphine are reduced in arrestin-3 knockout mice (39, 40).

Because one of the functions of dopamine receptors is to support the cell growth and proliferation required for neural plasticity, the downstream effectors that are activated by dopamine receptors include kinases involved in the regulation of these phenomena. There is particularly strong evidence that the pro-proliferation extracellular signal-regulated kinases 1/2 (ERK1/2) (41-43) and Src kinase (44) are among the key effectors regulated by dopamine signaling. Although these are well-studied effectors, there is an intense debate in the field as to how activated dopamine receptor selects between these pathways, and even debate as to whether G proteins or arrestins are the primary intermediary (30, 45-48). In general, the role of arrestins and G proteins must be experimentally studied in each case, and the answers are likely to be different (22, 27, 49)

Here, we used the D1R as a model receptor to explore global questions about the mechanisms of GPCR signal bias using ERK1/2 and Src kinase as model effectors. We first determined which aspects of dopamine-dependent ERK1/2 and Src signaling were mediated by arrestin versus G proteins. We further evaluated the effect of differential D1R phosphorylation on coupling to G protein, arrestin-3, and signal bias. Our results suggest complex input/output relationship in GPCR signaling.

## Results

### D1R activation of ERK1/2 and Src requires arrestin-3

Given the existing controversy in the field as to whether G proteins or arrestins mediate GPCR signaling to protein kinases, we first tested to what extent D1R-mediated ERK1/2 and Src activation requires G protein versus arrestin, focusing on Gαs and arrestin-3, which are characterized cognate signaling partners of D1R. To this end, we used two HEK293A cell lines. One lacked endogenous Gαs/Gα-olf, but retained all Gβγs and arrestins. The other lacked arrestin-2/3 but retained Gαβγ subunits (**Fig. 1A**). In these cells we assessed how arrestin-3 and Gαs affect dopamine-stimulated ERK1/2 and Src phosphorylation (**Fig. 1B-D**). We found that the addition of dopamine to HEK293 cells expressing wild-type (WT) D1R activates both ERK1/2 and Src in the absence of Gαs/Gα-olf **(Fig. 1B (left panel), 1C and 1D)**. In contrast, we did not observe significant dopamine-induced ERK1/2 or Src phosphorylation in HEK293 cells lacking the non-visual arrestins **(Fig 1B-D)**. This suggests that endogenous Gαs/Gα-olf (or other G protein subunits present in the arrestin-2/3 KO cells) cannot activate ERK1/2 or Src in the absence of arrestin-2/3. Or to put it another way, these data indicate that arrestin, and not Gαs/Golf, is necessary for D1R activation of both ERK1/2 and Src kinase.

**Figure 1.**
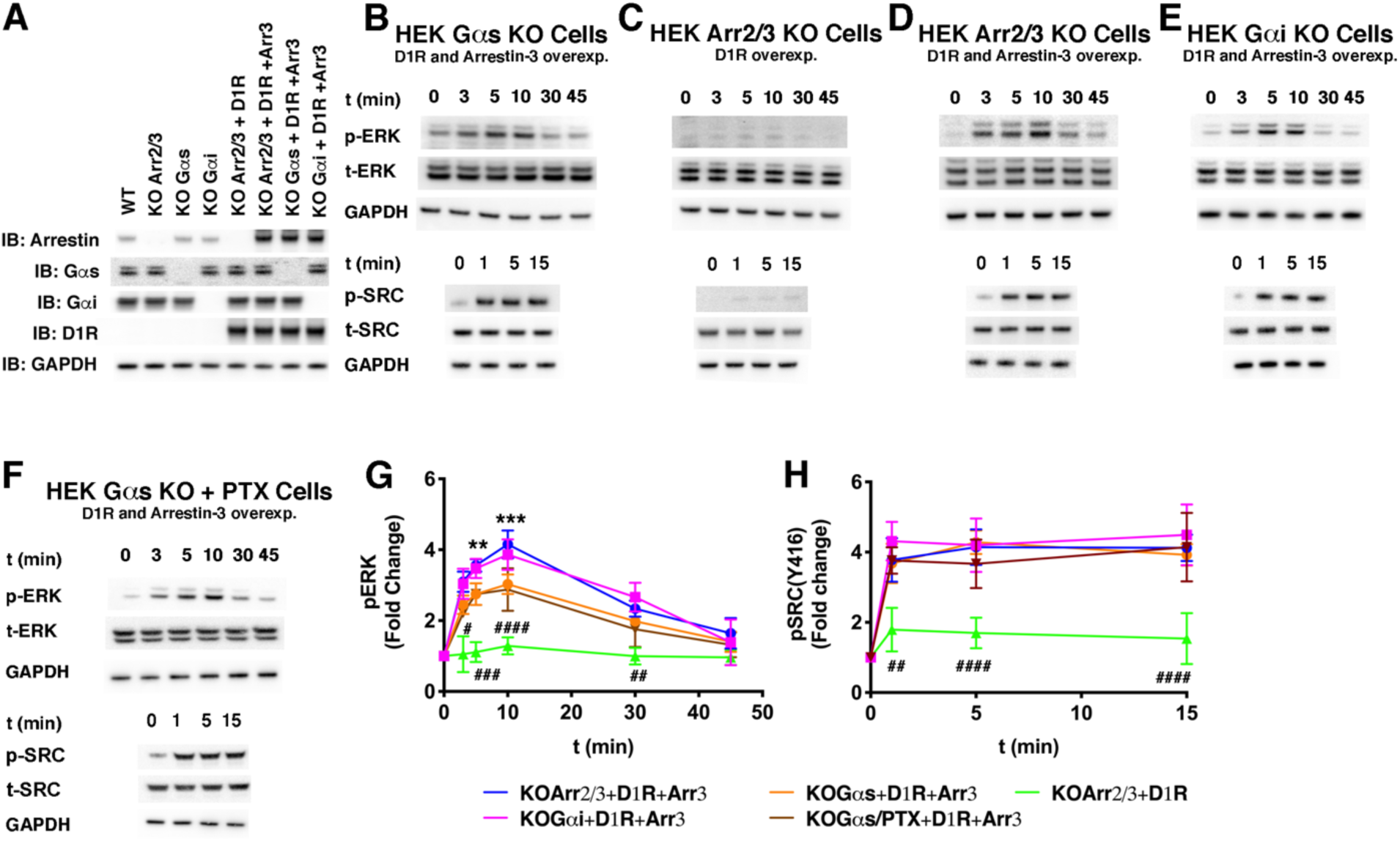
Effect of Gαs and arrestin-3 on D1R-dependent ERK1/2 and Src activation. (A) Indicated proteins were transiently expressed in wild-type or KO HEK293 cells. The arrestin antibody recognizes both arrestin-2 and arrestin-3. GAPDH was used as a loading control. (B - F) Time course of ERK1/2 and Src activation as monitored by immunoblotting of phosphorylated ERK1/2 (pERK1/2), phosphorylated Src (pSrc), total ERK1/2 (tERK1/2), and total Src (tSrc). Assays were initiated by adding 10 µM dopamine at 37 °C and performed in (B) Gαs knockout cells (C) arrestin-2/3 knockout cells, (D) arrestin-2/3 knockout cells with arrestin-3 reintroduced, (E) Gαi/o/t knockout cells expressing additional arrestin-3 or (F) Gαs knockout cells expressing both PTX (91) **(SI Appendix Fig. S1)** and additional arrestin-3. Gels are representative of at least three independent experiments. (G-H) Gels in panels (B) – (F) were quantified by densitometry and (G) ERK1/2 and (H) Src activation were expressed as the fold change relative to unstimulated cells. Orange trace, co-expression of D1R and arrestin-3 in Gαs/Gα-olf knockout cells; magenta trace, co-expression of D1R and arrestin-3 in Gαi/Gαo/Gαt knockout cells; green trace, ERK1/2 or Src activation in arrestin-2/3 knockout cells expressing D1R; blue trace, co-expression of D1R and arrestin-3 in arrestin-2/3 knockout cells. Statistical analysis was performed using one-way ANOVA followed by Tukey’s post hoc test with correction for multiple comparisons between D1R^WT^ mediated signal at each time point. In panel E, the significance shown as “*” is the comparison between “orange” and “blue” traces (**, p<0.01; ***, p<0.001) and “#” is the comparison between “blue” and “green” traces (#, p<0.05; ##, p<0.01; ###, p<0.001; ####, p<0.0001). In panel F, the significance was detected at every time point between “green” and “orange” (and “blue”). We didn’t detect significant differences between blue and magenta and also orange and brown traces in case of ERK and Src activation. Means +/- SD from at least three independent experiments are shown.

### G proteins enhance arrestin-dependent D1R activation of ERK1/2

We next tested whether there was G protein-dependent cross-talk in the system. To do this, we co-expressed D1R and arrestin-3 in arrestin-2/3 KO cells **(Fig. 1B - 1D)** and compared the activation of ERK1/2 and Src to the activation observed in the Gαs/Gα-olf KO cells **(Fig. 1B - 1D)**. This allowed us to assess the effect of Gs/G-olf on signaling because each of these cell lines has an equivalent amount of arrestin-3 (**Fig. 1A**). We did not detect a contribution of G protein to Src activation. Thus, arrestin-3 is both necessary and sufficient for full Src activation by dopamine-stimulated D1R. In contrast, we found that G protein-containing cells exhibited increased ERK1/2 activation when compared to Gαs/Gα-olf KO cells. Even though D1R primarily interacts with Gαs, GPCRs often display coupling promiscuity, i.e. they can interact with other G proteins. For instance, the β_2_AR can switch from Gαs to Gαi after phosphorylation by PKA (50). To test this possibility, we either co-expressed D1R and arrestin-3 in Gαi/o/t KO cells **(Fig. 1A and E)** or we co-expressed D1R, arrestin-3, and pertussis toxin (PTX) in Gαs/Gα-olf KO **(Fig. 1F, SI Appendix Fig S1)**. In both cases, we detected similar time-dependent ERK1/2 and Src phosphorylation as in arrestin-2/3 KO cells expressing D1R and arrestin-3. This result indicates that the increased ERK1/2 phosphorylation in cells expressing both G protein and arrestin-3 does not originate from D1R switching from Gαs to Gαi. Taken together, these data suggest Gαs likely cooperates with arrestin-3 in D1R-mediated ERK1/2 activation.

### Arrestin-3 binds specific phosphomimetic-containing peptides derived from the C-terminus and ICL3 of D1R

We next sought to elucidate the underlying mechanism, both in how arrestin activates the evolutionarily unrelated ERK1/2 and Src kinases and in how G protein enhances arrestin-dependent ERK1/2 activation. The simplest conceivable model is that arrestin binds to a phosphorylated receptor and to an effector in a signaling complex. The “barcode” hypothesis (51-53) implies that there are multiple conformations of active arrestin that are energetically similar, but that distinct receptor phosphorylation patterns increase the probability of arrestin adopting a particular active conformation. This in turn allosterically induces the formation of a binding site that prefers particular effectors. To test whether different phosphorylation patterns of receptor promote the binding of different effectors, we first identified the locations on D1R that interact with arrestin using peptide array analysis. We focused on two regions of D1R that contain a disproportionate number of S/T residues and that are exposed to the cellular interior: the C-terminus and ICL3.

The role of the fifteen S/T residues of the C-terminal region of D1R has previously been investigated by truncation mutagenesis (38), suggesting a model where this region is not the primary arrestin binding site, but instead shields the primary binding site in ICL3. Here, we focused on a patch of three S/T residues near the distal C-terminus to investigate whether arrestin-3 could bind directly to this region. Two 15-mer peptides, D1R^F371-E385^ **(SI Appendix Fig. S2A**) and D1R^Y426-D440^ **(Fig. 2A)** corresponding to the distal region of human D1R C-terminus **(Fig. 2A)** contain either four (D1R^F371-E385^) or three (D1R^Y426-D440^) S/T residues that can be phosphorylated by GPCR kinases (GRKs). For peptides corresponding to D1R^F371-E385^, single phosphomimetic substitutions of S372 or S373 did not affect arrestin-3 binding, while those at S380 or S382 increased it (**SI Appendix Fig. S2B**). The effect of the S380 and S382 mutations was additive, however, the impact was not straightforward because the phosphomimetic substitution of S372 and S373 interfered with arrestin binding in compound mutants, i.e. the quadruple mutant had lower arrestin-3 binding than the S380/S382 double mutant and similar arrestin-3 binding as the individual S380 and S382 single mutants (**SI Appendix Fig. S2B**). These data suggest that this site may not be relevant for arrestin-dependent signaling. In contrast, D1R^Y426-D440^ peptides with single substitutions of T428, S431 or T439 with phosphomimetics demonstrate a modest, but statistically significant, increase in binding to arrestin-3 **(Fig 2B)**. These effects were additive, with double mutants exhibiting greater binding than single mutants, and a triple mutant demonstrating the largest increase in arrestin-3 binding **(Fig 2B)**. The T428, S431 and T439 were previously shown to be phosphorylated after D1R activation and to be important for arrestin-mediated D1R internalization (54, 55). Peptides with alanine substitutions at these positions exhibited binding comparable to the native sequence (**Fig. 2B**), indicating that negative charges, rather than H-bonding capability, are necessary.

**Figure 2.**
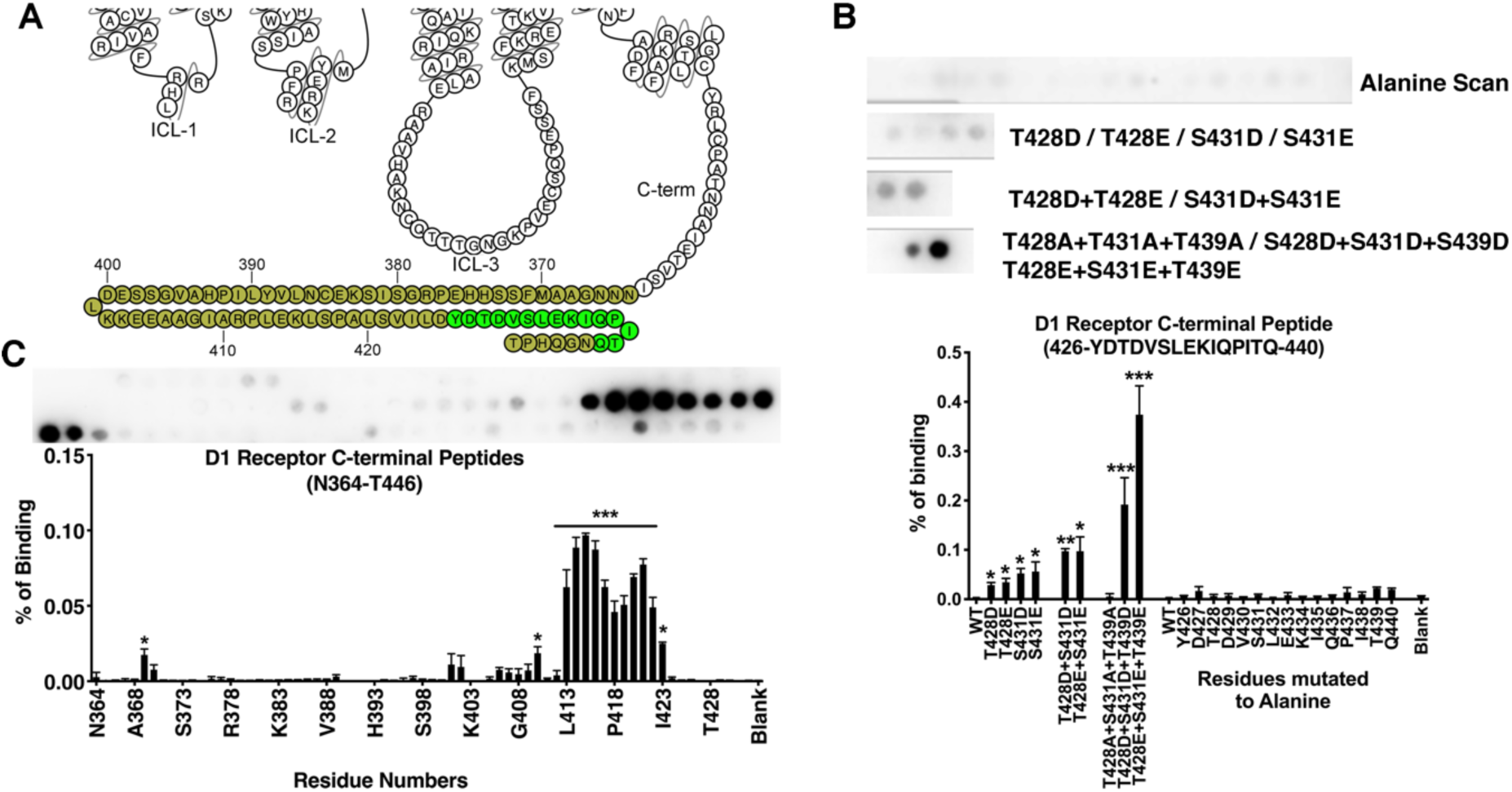
Peptide array analysis of the D1R C-terminus. (A) A snake diagram (92) highlighting the residues that were tested for their role in arrestin-3 binding. (B) Peptide array analysis of the D1R^426-440^ peptide. As monitored by Far Western analysis (see Methods), the wild-type peptide exhibited little detectable interaction with arrestin-3; alanine scanning did not significantly affect arrestin-3 binding. Phosphomimetic substitution at positions T228, S431, and T439 enhanced binding, the effect of these mutations was additive. (C) A scan of 15-mer peptides derived from the C-terminus. The residue in the lower graph indicate the N-terminal residue of each peptide (i.e. “N364” corresponds to the D1R^364-378^ peptide, with the sequence NNNGAAMFSSHHEPRG). Means +/- SD of at least three independent experiments are shown. Representative peptide array results are shown above graphs displaying averages. Statistical analysis was performed using one-way ANOVA followed by Tukey’s post-hoc test. The statistical significance of the difference of the signal from the WT peptide spot is shown, as follows: *, p<0.05; **, p<0.01; ***, p<0.001.

Interestingly, a peptide array scan of the entire native sequence of the D1R C-terminus (D1R^N364^ to D1R^T446^) in one amino acid shifts identified a short region (D1R^L413^ to D1R^I423^) immediately N-terminal to T428/S431/T439 that bound strongly to arrestin-3 without phosphomimetic substitutions **(Fig. 2C)**. All of the unphosphorylated peptides that exhibited a strong interaction with arrestin-3 contain a “DYD” sequence motif and a net negative charge. Conceivably, this region functions as a pre-docking site for arrestin-3 that allows a stable interaction with unphosphorylated receptor. This possibility is consistent with prior reports that some receptors, for example the D2 dopamine receptor and the M2 muscarinic receptor, interact robustly with arrestin even in the absence of receptor phosphorylation (56). An alternative possibility is that in the context of folded D1R in the membrane, this region of the C-terminus could be shielded and inaccessible for interacting proteins.

We next used peptide array analysis to evaluate how phosphomimetic substitution of S/T residues of ICL3 affects arrestin binding. Previously studies that focused on specific residues of D1R-ICL3 showed that phosphorylation of S258, S259 and T268 is important for receptor desensitization (38, 55). D1R-ICL3 contains only 29 amino acids. Seven of these residues are S/T, so a comprehensive combinatorial approach would result in 5,039 potential phosphorylation patterns. Therefore, we simplified our analysis by dividing this region into two 15-mer peptides (**Fig. 3**), equivalent to D1R^A238-E252^ and D1R^E252-R266^. For both peptides, the native sequence exhibited little measurable binding to arrestin-3. We then evaluated the impact of phosphomimetic substitution on arrestin-3 binding.

**Figure 3.**
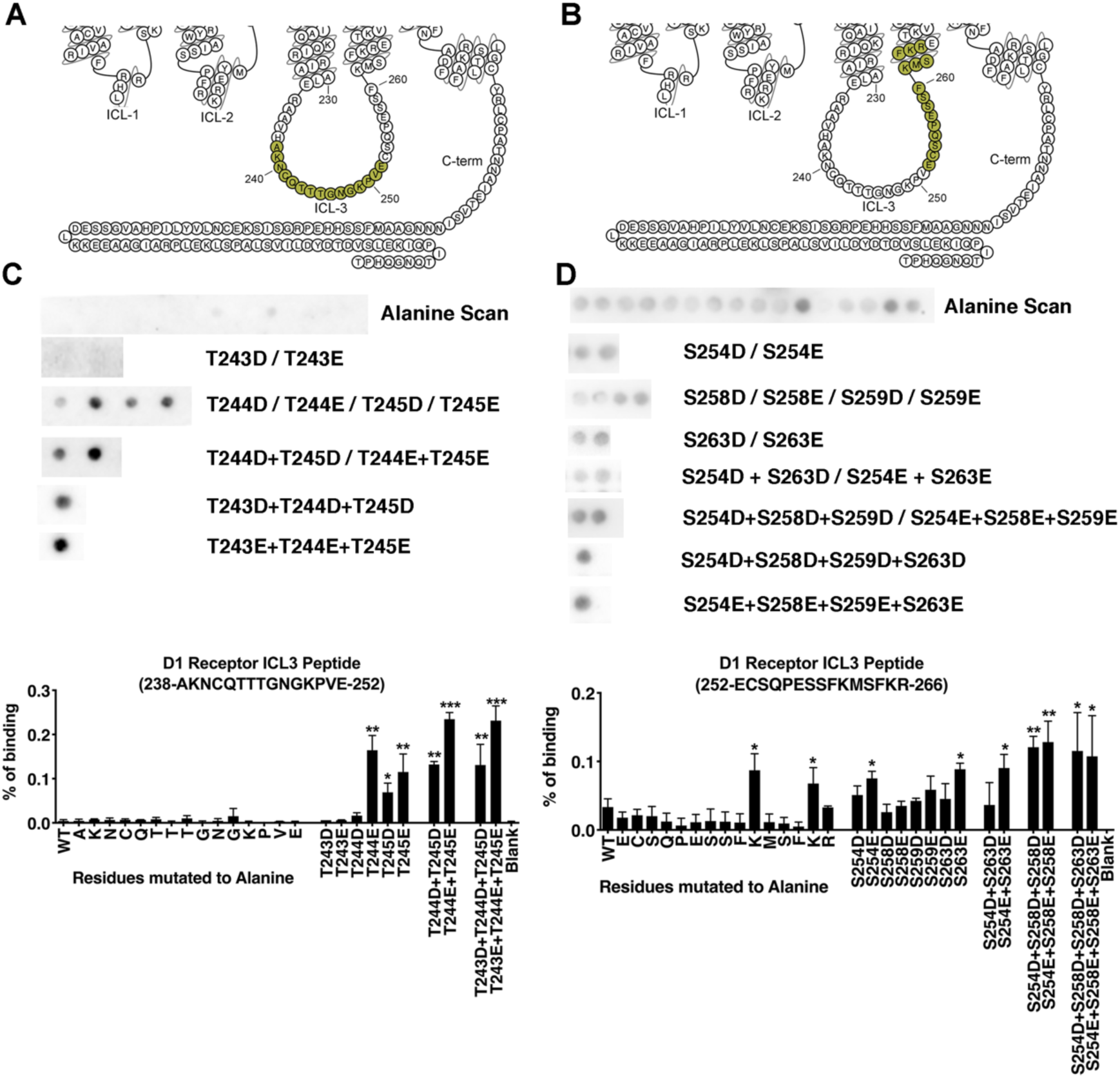
Peptide array analysis of the D1R ICL3. Peptide array analysis was used to identify residues in the D1R ICL3 that interact with arrestin-3. (A) – (B) snake diagrams showing the residues in tested ICL3-derived peptides (92). Representative Far Western analysis and quantitation of the D1R^238-252^ peptide (C) and D1R^252-266^ peptide (D) following alanine scanning and phosphomimetic substitutions. Means +/- SD of 3-5 independent experiments are shown. Statistical analysis was performed using one-way ANOVA followed by Tukey’s post-hoc test. The statistical significance of the difference of the signal from the WT peptide spot is *, p<0.05; **, p<0.01; ***, p<0.001.

The D1R^A238-E252^ peptide contains three adjacent threonines, T243, T244 and T245 **(Fig. 3A)**. Single or double phosphomimetic substitutions of T244 or T245 increased the binding of arrestin-3 **(Fig. 3A)**, while phosphomimetic substitution of T243 did not affect binding (**Fig. 3A**). In the D1R^A238-E252^ peptide, alanine scanning did not affect arrestin-3 binding.

The D1R^E252-R266^ peptide contains four putative phosphorylation sites: S254, S258, S259 and S263. Single phosphomimetic substitutions of S254 and S263 increased arrestin-3 binding, while substitutions of S258 and S259 did not. A triple substitution of S254, S258, S259 resulted in the highest binding of ICL3 to arrestin-3 (**Fig. 3B**). In the D1R^E252-R266^ peptide, alanine scanning identified two lysines where neutralization increased binding (**Fig. 3B**); both substitutions resulted in a net negative charge of the resultant peptide.

### Only some S/T residues of D1R contribute to the arrestin-3 recruitment to D1R in cells

We next tested which of the identified residues support the recruitment of arrestin-3 to D1R in HEK293 cells. Because of the complexity of the system, we used biased randomization to design single, double, triple, and quadruple alanine and phosphomimetic mutations **(SI Appendix Fig. 3)**. Using an in-cell BRET assay (**Fig. 4A**) (46), we evaluated which S/T residues decreased arrestin-3 binding to D1R when mutated to alanine. For a subset of the mutations that reduced binding upon alanine substitution, we tested whether mutation to a phosphomimetic increased arrestin-3 binding to D1R.

**Figure 4.**
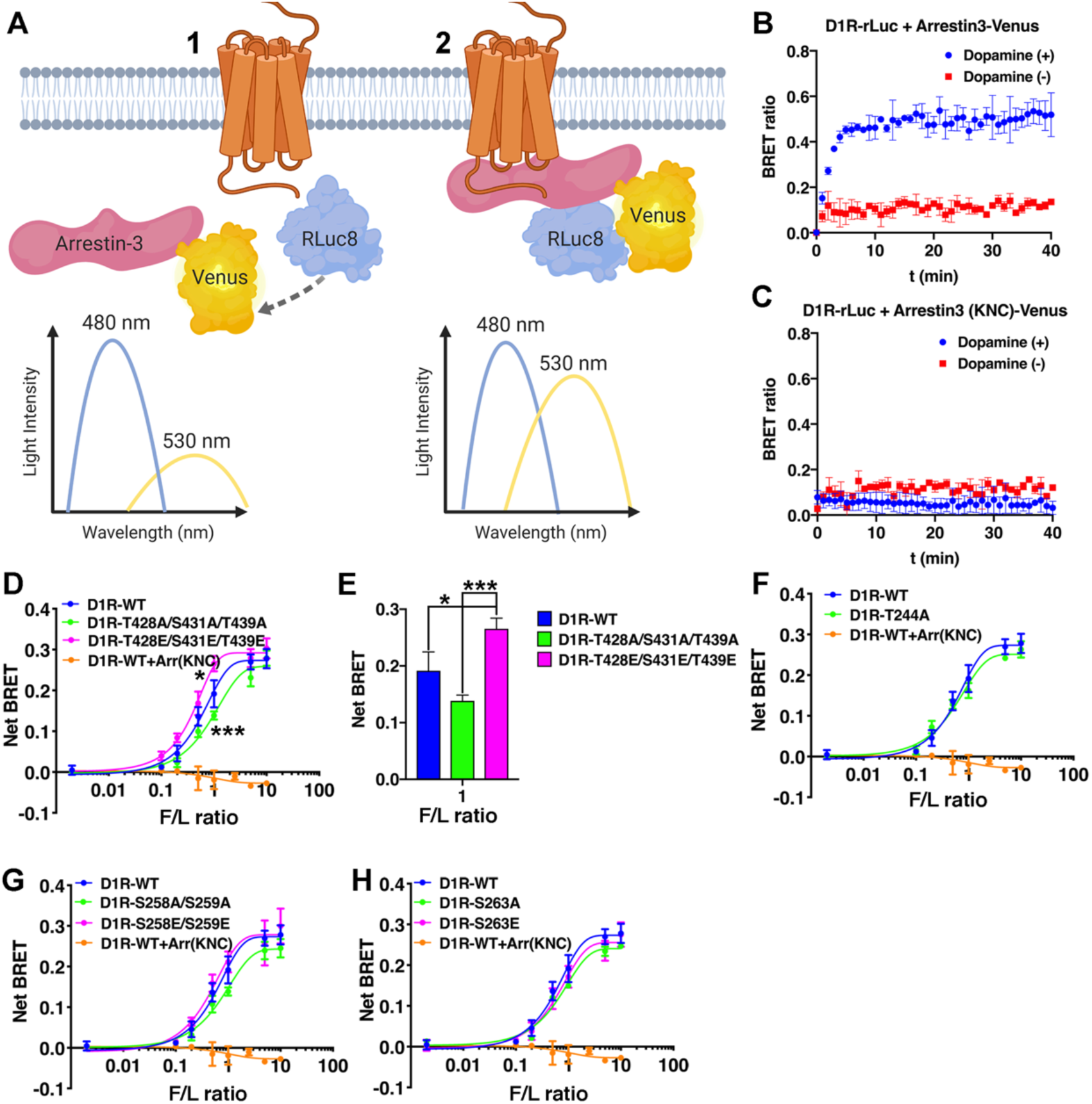
In-cell binding of arrestin-3 to WT and mutant D1R. (A) Schematic of the BRET assay used to measure binding between WT and mutant D1R-Rluc8 to Venus-arrestin-3 in arrestin-2/3 KO HEK293 cells. The maximum emission of Rluc8 is at 480 nm (1). When the two proteins (D1R and arrestin-3) are in close proximity (<10 nm), the energy is transferred to Venus, which emits light at 530 nm (2). Figure made using BioRender (BioRender.com) (B) Raw BRET signal between WT-D1R and WT-arrestin-3 (1:5 donor-acceptor ratio) in the presence (blue trace) or absence (red trace) of 10 µM dopamine. (C) Raw BRET signaling between WT-D1R and the Arr3^KNC^ negative control. (D) BRET signal of wild-type and D1R C-terminal mutant receptor stimulated by 10 µM dopamine in the presence of increasing amounts of Venus-arrestin-3 (0-1µg) and monitored for 10 min. The net BRET ratio was calculated as the acceptor emission divided by the donor emission and expressed as the change from unstimulated cells. (E) The statistical significance of the signal detected between the WT-D1R and the D1R-T428E-S431E-T439E (*, p<0.05) and between the D1R-T428A-S431A-T439A and the D1R-T428E-S431E-T439E (***, p<0.001) in 1:1 F/L ratio. (F) – (H) BRET signal of wild-type and D1R ICL3 mutant receptors. **SI Appendix Fig. S4** shows BRET data collected from all receptor mutants. Means +/- SD of 3-5 independent experiments are shown. Statistical analysis was performed by one-way ANOVA followed by Tukey’s post-hoc test.

Dopamine stimulation of arrestin-2/3 KO HEK293 cells co-expressing Venus-arrestin-3 with Luciferase-tagged wild-type D1R (RLuc8-D1R^WT^) resulted in a robust, agonist-dependent BRET signal (**Fig. 4B**) that saturates with increasing Venus-arrestin-3 expression (57-59). Conversely, we did not observe a detectable BRET signal following dopamine stimulation of arrestin-2/3 KO HEK293 cells co-expressing RLuc8-D1R^WT^ and the established negative control arrestin-3^KNC^ (60), a mutant arrestin-3 that does not bind GPCRs (**Fig. 4C**).

Simultaneous substitution of the three C-terminal S/T residues of D1R with alanine (D1R^T428A/S431A/T439A^) modestly decreased arrestin-3 recruitment, while phosphomimetic substitution at the same positions (D1R^T428E/S431E/T439E^) modestly increased it (**Fig. 4D-E**). This suggests that phosphorylation of these C-terminal residues contributes to the recruitment of arrestin-3 to activated D1R. Conversely, all mutants within ICL3 showed a BRET signal statistically indistinguishable from that observed for the D1R^WT^-RLuc8 **(Fig. 4F-H, SI Appendix Fig. S4)**. Although, the peptide array analysis indicated that phosphomimetic-containing peptides corresponding to ICL3 bind arrestin-3 *in vitro*, of the putative phosphorylation sites tested here, only those at the C-terminus appear to contribute significantly to arrestin-3 recruitment to D1R in cells. It is clear that arrestin-3 is still recruited when this C-terminal region is altered, suggesting that other regions of D1R contribute to arrestin-3 binding. The negatively charged residues within the D1R^L413-I423^ region, identified by the peptide array analysis (**Fig. 2C**) are likely contributors to the arrestin-3-D1R interaction. One interpretation of the peptide array and BRET results is that the sites in ICL3 are important for directing the function of arrestin following receptor binding.

Previous studies by the Sibley group suggest a model where the ICL3 is the true arrestin binding site in D1R (38). In this model, the unphosphorylated C-terminus of D1R shields ICL3 and phosphorylation of the C-terminus elicits a conformational change that displaces the C-terminus and exposes ICL3 (38, 55). In this context, our results suggest that initial recruitment of arrestin to receptors involves a different binding mode than is observed in the final arrestin-receptor signaling complex, an idea originally discussed by Shukla *et*.*al* (61, 62). Thus, it is possible that the sites that we tested in the D1R ICL3 do not appear to be significant for the initial recruitment of arrestin-3 to receptor because the mutagenesis did not promote the conformational change suggested by the Sibley group (38). Alternatively, ICL3 may be important for directing the function of arrestin-3 even though it is not involved in the initial recruitment of arrestins.

### Differential effect of the D1R phosphorylation pattern on ERK1/2 and Src activation

We selected ERK1/2 and Src as model effectors to understand how the phosphorylation pattern of D1R directs function. We evaluated the time course of ERK1/2 and Src activation in arrestin-2/3 K/O HEK293 cells co-expressing C-terminally HA-tagged arrestin-3 (arrestin-3^HA^) and either wild-type D1R (D1R^WT^) or D1R variants containing alanine or phosphomimetic substitutions of critical S/T residues **(Fig. 5)**.

**Figure 5.**
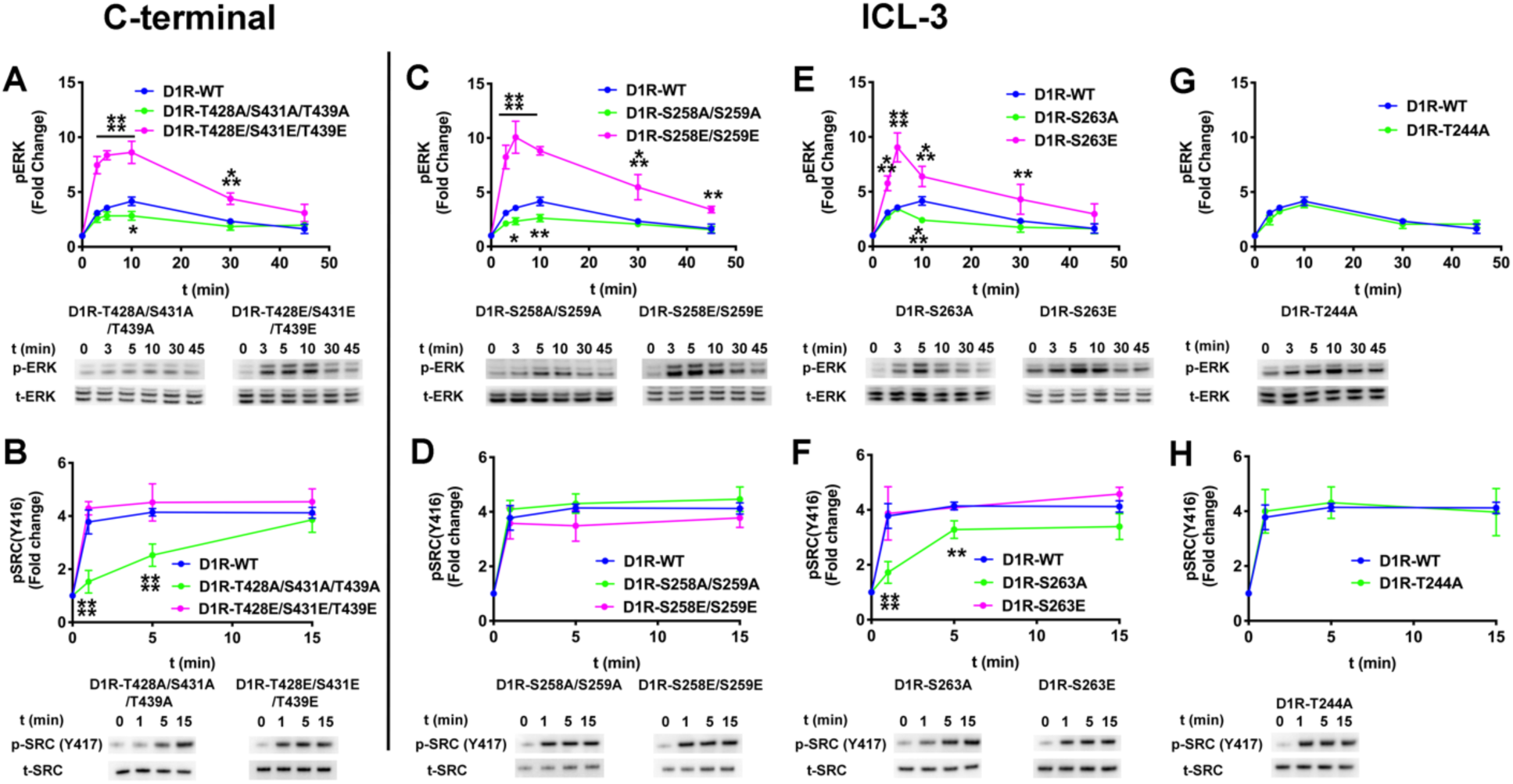
The effect of the D1R-ICL3 phosphorylation pattern on ERK1/2 and Src phosphorylation. Arrestin-2/3 KO HEK-293 cells were co-transfected with plasmids encoding arretsin-3HA and wild-type or mutant D1R. The assay was initiated by adding 10 µM dopamine at 37 °C. Cell lysates were separated on a 4-12% gradient SDS-PAGE, transferred to PVDF membranes and immunoblotted for phosphorylated ERK1/2 (pERK), phosphorylated Src (pSrc), total ERK1/2 (tERK), and total Src (tSRC). Signals were quantified by densitometry and expressed as the fold change relative to unstimulated cells. (A, C, E, G) Time course of ERK1/2 phosphorylation upon dopamine stimulation. (B, D, F, H) Time course of Src activation upon dopamine stimulation. Means +/- SD of at least three independent experiments are shown. D1R, Arrestin-3 and GAPDH expression levels are shown in the **SI Appendix Fig. S7**. GAPDH expression was used as a loading control. Statistical analysis was performed with one-way ANOVA followed by Tukey’s post hoc test with correction for multiple comparisons between wild-type and mutant D1R mediated signal at each time point. Significance is shown, as follows: *, p<0.05; **, p<0.01; ***, p<0.001; ****, p<0.0001.

As compared to D1R^WT^, the D1R^T428A/S431A/T439A^ mutant elicited lower ERK1/2 activation up to 10 min, and a marked slowing of Src activation. Phosphomimetic substitutions at the same locations (D1R^T428E/S431E/T439E^) increased ERK1/2 activation at all timepoints but did not affect Src activation **(Fig. 5A and B)**. The reduced activation of both ERK1/2 and Src by the triple alanine mutant D1R^T428A/S431A/T439A^ is consistent with these residues contributing to receptor recruitment of arrestin-3 (**Fig. 4E**). The difference in ERK1/2 activation as compared to Src suggests an additional role of these residues in directing the signal.

The mutations of the D1R ICL3 yielded three distinct signaling outcomes. The first is exemplified by any form of D1R containing mutations of S258 and S259. D1R^S258A/S259A^ induced significantly lower ERK1/2 phosphorylation than WT at 5- and 10-min., but did not affect Src activation **(Fig. 5C,D, SI Appendix Fig. S5,6)**. The phosphomimetic mutations in D1R^S258E/S259E^ significantly enhanced ERK1/2 activation at all time points **(Fig. 5C, magenta)** without affecting Src activation. The data suggest that phosphorylation of S258 and S259 in D1R directs the signaling via arrestin-3 toward ERK1/2 activation.

The second outcome was observed with mutations of S263, T244/S263, and T245/S263 and appeared similar to what we found for the C-terminal mutants. The D1R^S263A^, D1R^T244A/S263A^, and D1R^T245A/S263A^ mutants elicited slowed activation kinetics of both ERK1/2 and Src than D1R^WT^ **(Fig. 5E,F, SI Appendix Fig. S5,6)**. Specifically, ERK1/2 activation was lowered at the 10-min time point **(Fig. 5E)** and Src **(Fig. 5F)** activation was significantly less than with D1R^WT^ at early time points, with later activation comparable to that induced by D1R^WT^. The introduction of phosphomimetics at these positions enhanced ERK1/2 activation above the levels of D1R^WT^ **(Fig. 5E, magenta)**. These results identify S263 as important for the kinetics of activation of both pathways. They also identify phosphorylation of S263 as important for directing the signal toward the activation of ERK1/2.

The third outcome was observed with the D1R^T244A^, D1R^245A^, and D1R^244A/245A^, which induced ERK1/2 and Src phosphorylation at levels similar to the D1R^WT^ **(Fig. 5G and H, SI Appendix Fig. S5,6)**. Thus, while the peptides containing phosphomimetics at T244 and T245 affect arrestin-3 binding in vitro (**Fig. 3**), these residues do not appear to be involved in the initiation of either ERK1/2 or Src signaling. As these also do not affect binding of arrestin-3 to the receptor in cells (**Fig. 4F**), it is possible that they are not accessible to arrestins in the context of the folded receptor. Alternatively, the phosphorylation of these residues could direct signaling toward pathway(s) that we did not investigate here.

We also tested the effect of a possible G protein switch (from Gαs to Gαi) during D1R activation on ERK and SRC activation. We evaluated ERK1/2 and Src activation in Gαi/o/t KO HEK293 cells co-expressing arrestin-3HA and either D1R^WT^ or D1R mutants (**SI Appendix Fig. S8**). Our data showed similar ERK and Src phosphorylation patterns with arrestin-2/3 K/O cells which co-expressed D1R and arrestin-3. This result suggests that Gαi proteins do not play a significant role in ERK1/2 and Src activation.

### Mechanism of Gs enhancement of the arrestin-dependent activation of ERK1/2

With a better understanding of how phosphorylation pattern directs arrestin-dependent signaling toward ERK1/2 or Src, we next wanted to understand how Gs cooperates with arrestin in activation of ERK1/2, but not Src **(Fig. 1)**. There are multiple mechanisms by which this could occur. <u>First</u>, the D1R-arrestin-3-ERK1/2 signaling complex could directly incorporate Gs into a super-complex, as was described for a chimeric β2-adrenergic receptor containing the V2 vasopressin receptor C-terminus (63, 64). <u>Second</u>, Gs could compete with arrestin-3 for binding to active D1R, resulting in a mixed population of D1R-Gs and D1R-arrestin-3 complexes with cross-talk enhancing ERK1/2 activation. <u>Third</u>, dopamine-independent Gs activation could enhance arrestin-dependent ERK1/2 activation via cross-talk that does not require continued Gs activation.

To determine whether a D1R-arrestin-3-Gs-ERK1/2 complex forms in cells, we performed a BRET assay using Venus-tagged arrestin-3 and eYFP-tagged Gs. We did not observe statistically significant BRET signal (**SI Appendix Fig. S9**) under our experimental conditions. This suggests that no supercomplex is formed, but does not exclude the possibility of a D1R-arrestin3-Gs complex that is not detectable by this method.

To determine whether a population of D1R-arrestin-3 and D1R-Gs was present in cells, we monitored in-cell Gs recruitment to D1R^WT^ and D1R phosphomimetic variants in the absence and presence of arrestin-3. In arrestin-2/3 KO cells, we found that phosphomimetics on D1R did not directly impact Gs recruitment (**SI Appendix Fig. S10)** suggesting that receptor phosphorylation in and of itself does not terminate G protein coupling. However, in cells containing arrestin, Gs coupling is significantly reduced when D1R contains phosphomimetics in the ICL3 (D1R^S258E/ S259E^, and D1R^S263E^), but not at the C-terminus (D1R^T428/S431/T439^) (**SI Appendix Fig. S11)**. This is consistent with a competition between arrestin-3 and Gs and a co-existence of D1R-arrestin-3 and D1R-Gs complexes.

To evaluate D1R-independent cross-talk at another point in signaling, we tested the impact of Gs on effector activation by phosphorylation-deficient and phosphomimetic-containing D1R mutants. If the influence of Gs on arrestin-3-dependent ERK1/2 activation is indirect, we expected that Gs would enhance ERK1/2 activation by these variants. We therefore selected four pairs of D1R mutants (D1R^T244A^/D1R^T244E^, D1R^S258A/S259A^/D1R^S258E/S259E^, D1R^S263A^/D1R^S263E^ and D1R^T428A/S431A/T439A^/D1R^T428E/S431E/T439E^) representing different signaling outcomes and measured ERK1/2 and Src activation in Gs KO HEK293 cells **(Fig. 6)**. All four alanine mutants (i.e. D1R^T244A^, D1R^S258A/S259A^, D1R^S263A^, and D1R^T428A/S431A/T439A^) showed a similar time course of both ERK1/2 and Src phosphorylation in the absence of Gs **(Fig. 6)** as they did in the presence of Gs (**Fig. 5**). In contrast, the phosphomimetic mutants (i.e. D1R^T244E^, D1R^S258E/S259E^, D1R^S263E^, and D1R^T428E/S431E/T439E^) showed an attenuated ERK1/2 activation in the absence of Gs **(Fig. 6A,C,E,D)**, as compared to cells expressing Gs (**Fig. 5A,C,E,D**). Interestingly, the D1R^T428E/S431E/T439E^ mutant yielded a slight increase in Src activation in the absence of Gs (**Fig. 6B, 4H**), while Src activation by the other phosphomimetic variants was unaffected by the presence of Gs (compare **Fig. 6B,D,F,H** with **Fig. 5B,D,F,H**).

**Figure 6.**
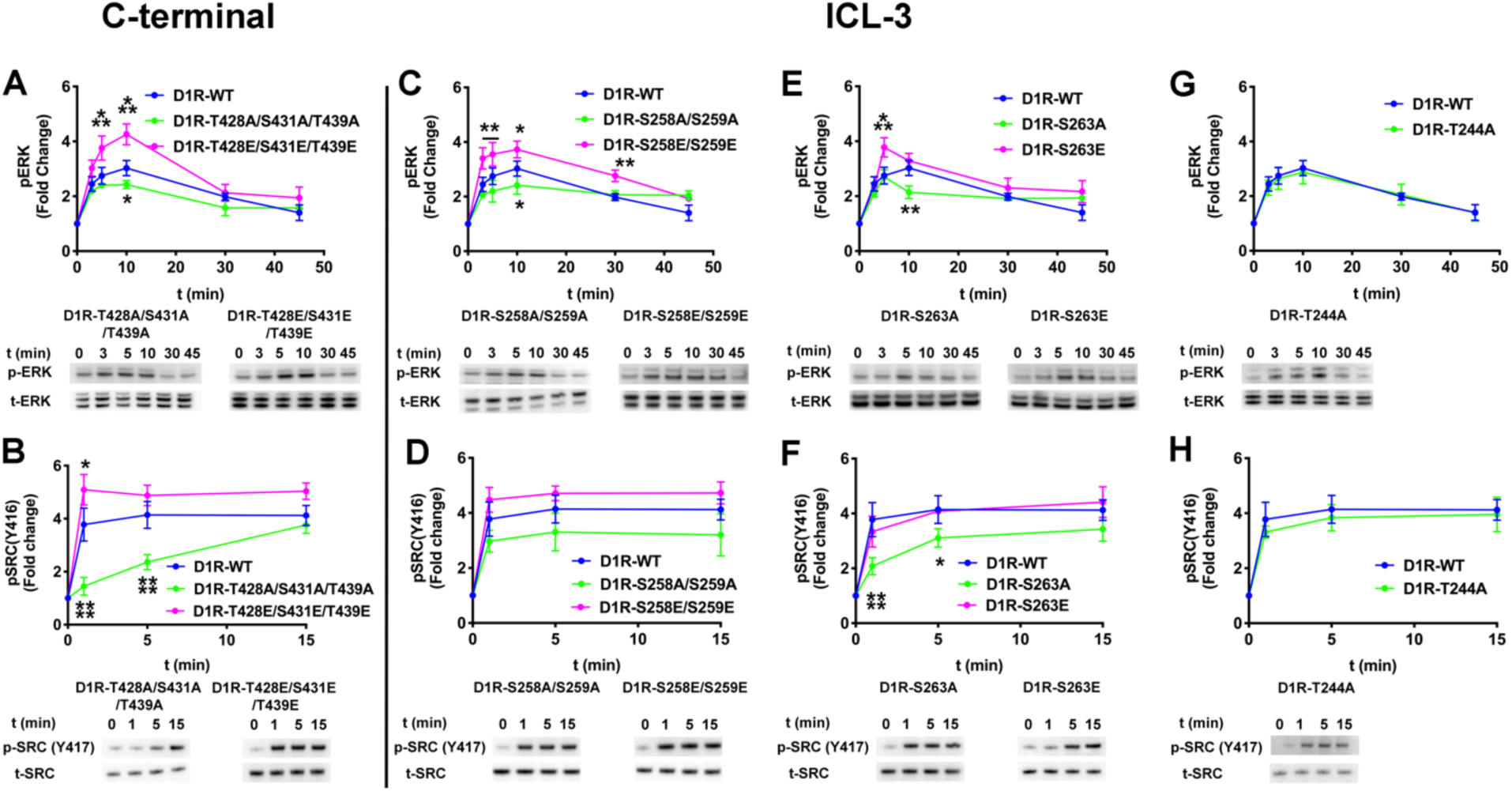
The effect of Gαs on arrestin-dependent ERK1/2 and Src activation by D1R mutants. Gαs/Gα-olf KO HEK-293 cells were co-transfected with plasmids encoding arretsin-3HA and indicated variant of D1R. The assay was initiated by 10 µM dopamine at 37 °C. Phosphorylated ERK1/2 (pERK), phosphorylated Src (pSrc), total ERK1/2 (tERK), and Src (tSrc) were visualized by Western blotting. Signals were quantified by densitometry and expressed as the fold change relative to unstimulated cells. (A, C, E, G) Time course of ERK1/2 phosphorylation upon dopamine stimulation. (B, D, F, H) Time course of Src activation upon dopamine stimulation. Means +/- SD of at least three independent experiments are shown. D1R, Arrestin-3 and GAPDH expression levels are shown in the **SI Appendix Fig. S7**. GAPDH expression was used as a loading control. Statistical analysis was performed with one-way ANOVA followed by Tukey’s post hoc test with correction for multiple comparisons between wild-type and mutant D1R mediated signal at each time point. Significance is shown, as follows: *, p<0.05; **, p<0.01; ***, p<0.001; ****, p<0.0001.

We then monitored the contribution of Gs to ERK1/2 and Src activation in arrestin-2/3 KO cells containing the same D1R mutants (**Fig. 7**). Surprisingly, we detected arrestin-independent ERK1/2 phosphorylation induced by D1R containing phosphomimetics in the C-terminus (i.e. D1R^T428E/S431E/T439E^), albeit at a level lower than the arrestin-dependent signaling (**Fig. 1, 5, 6)**.

**Figure 7.**
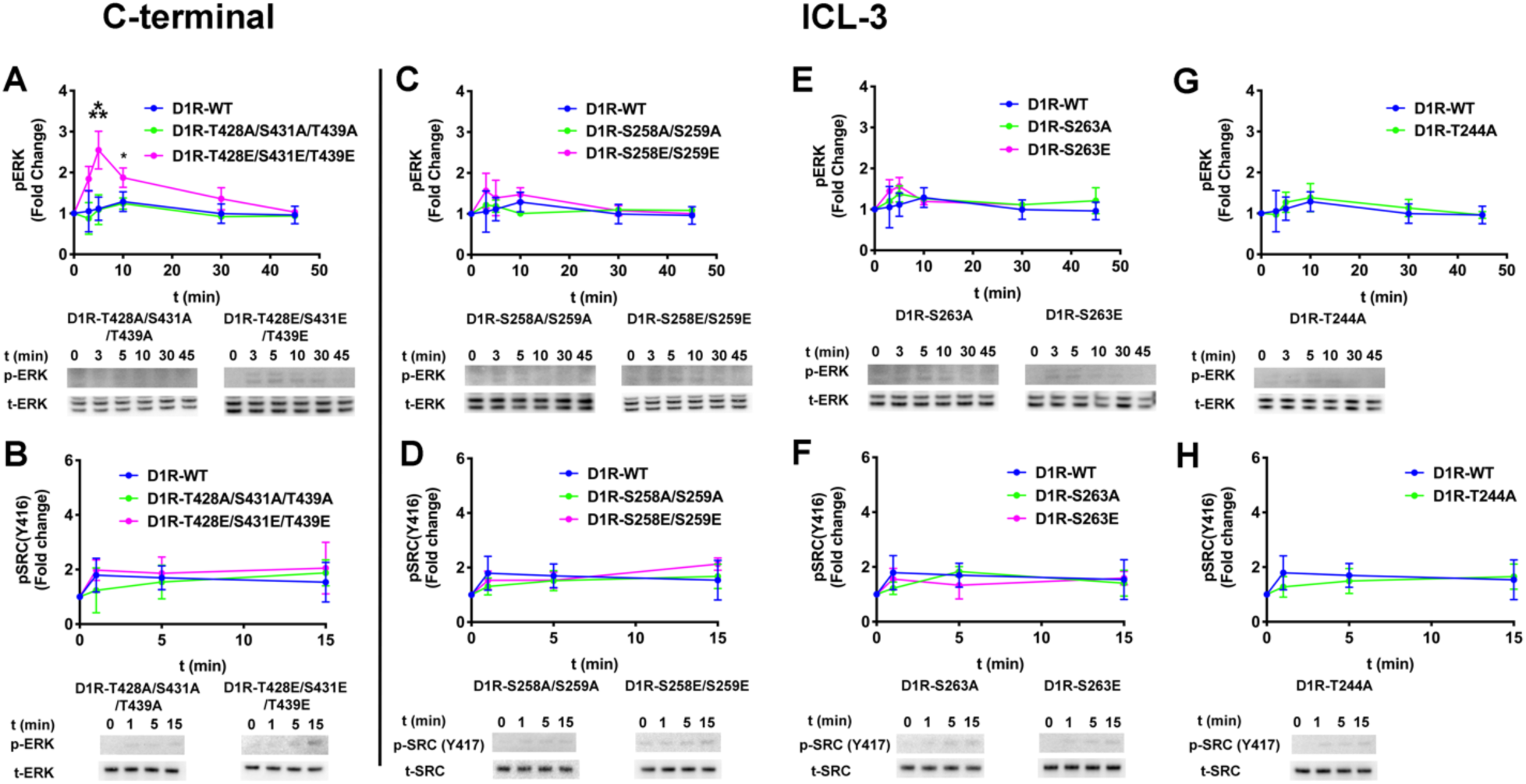
Gαs-dependent ERK1/2 and Src activation by D1R mutants. Arrestin-2/3 KO HEK-293 cells were co-transfected with plasmids encoding wild-type or mutant D1R. The assay was initiated by 10 µM dopamine at 37 °C. Cell lysates were separated on a 4-12% gradient SDS-PAGE, transferred to PVDF membranes, and immunoblotted for phosphorylated ERK1/2 (pERK), phosphorylated Src (pSrc), total ERK1/2 (tERK), and total Src (tSRC). Signals were quantified by densitometry and expressed as the fold change relative to unstimulated cells. (A, C, E, G) Time course of ERK1/2 phosphorylation upon addition of 10 μM dopamine. (B, D, F, H) Time course of Src activation upon dopamine stimulation. Means +/- SD of at least three independent experiments are shown. D1R and GAPDH expression levels are shown in the **SI Appendix Fig. S7**. GAPDH expression was used as a loading control. Statistical analysis was performed with one-way ANOVA followed by Tukey’s post hoc test with correction for multiple comparisons between wild-type and mutant D1R mediated signal at each time point. Significance is shown, as follows: *, p<0.05; **, p<0.01; ***, p<0.001; ****, p<0.0001.

## Discussion

Cells use a number of mechanisms to encode information. Signal transduction is amongst the most complex biological coding problems, and one where the code has not yet been cracked. Receptors interact with multiple types of ligands, with many factors fine-tuning the downstream signaling. Here, we evaluated elements of the signaling code that could explain how dopamine stimulation of D1R directs signaling toward ERK1/2 versus Src activation.

The relative roles of G protein and arrestin in ERK1/2 activation are hotly debated (22, 45-47). Our data clearly indicate that arrestin-3 is sufficient for both ERK1/2 and Src activation downstream of D1R (**Fig. 1**). This is in stark contrast with what has been recently reported for several other GPCRs, where ERK1/2 activation was found to depend on G proteins (30, 45-48). Our data further indicate that part of the biasing mechanism results from the receptor phosphorylation pattern, i.e. the phospho-barcode, affecting both arrestin-3 and Gαs coupling (**SI Appendix Fig. S11**).

A model for how this would occur is based upon the hypothesis that receptor-arrestin and receptor-G protein complexes coexist in a population; here, different phosphorylation patterns might result in different affinities for arrestin, with high affinity patterns allowing arrestin to compete effectively against G protein and lower affinity patterns resulting in co-existence of receptor-G protein and receptor-arrestin complexes in the cell. Conceptually, a framework for this model is extrapolated from findings on the well-studied rhodopsin and arrestin-1. In rhodopsin, lack of phosphorylation results in abnormally prolonged G protein-dependent signaling because arrestin does not compete with G protein for receptor binding (65). Moreover, biochemical data show that the receptor-arrestin interaction is enhanced more than 10-fold when the receptor is both phosphorylated and activated, as compared to receptor that is only activated or only phosphorylated. Thus, only simultaneous activation and phosphorylation of the receptor allows arrestin to outcompete G protein (66).

Crystal structures of the receptor-arrestin complexes indeed reveal that arrestin binds to both the phosphorylated elements in the C-terminus and the pocket unique to active receptor (the main G protein binding site)(67-70). While phosphorylation commonly reduces G protein coupling (71-73), in some cases it changes the G protein subtype that couples to receptor (50, 74). Phosphorylation per se does not preclude G protein activation, and our data show that phosphomimetics in the D1R do not significantly impact Gαs coupling (**SI Appendix Fig. S10 and 11**).

In contrast, arrestin binding to active phosphorylated GPCRs prevents G protein coupling via a direct competition (reviewed in (75)). Indeed, our data show that different phosphomimetic patterns on D1R affect Gαs coupling only in the presence of arrestin-3, which suggests co-existence of D1R-Gs and D1R-arrestin-3 complexes in the cell. Intriguingly, the effect of the phosphorylation barcode on arrestin-3 coupling means that the barcode also affects the likelihood of Gαs coupling, albeit indirectly. This model is consistent with our finding that ERK1/2 activation was enhanced by the presence of Gαs while Src activation is not affected by G protein. A synergistic effect of G proteins and arrestins is contrary to the classical paradigm of G protein signaling, but has been reported for growth hormone secretagogue receptor, neurokinin-1 receptor (76-79), and 5-HT_1B_-mediated activation of ERK1/2 (80). We propose that the synergistic effect is via co-existing populations of receptors, some coupled to arrestin and some coupled to G protein.

In addition to identifying how different phosphorylation positions of D1R influence whether signaling proceeds through arrestin-3, Gs, or both, these studies show how the barcode directs arrestin-dependent signaling to different effectors. Prior studies have shown that arrestins require at least three receptor-attached phosphates for high-affinity binding (67, 81-83), whereas most GPCRs have many more potential phosphorylation sites. In some cases, the phosphorylation of the primary sites in receptor permits the phosphorylation of additional residues (84). This type of hierarchical phosphorylation has been described for several GPCRs, including D1R (38), A3 adenosine receptor (85), and the δ-opioid receptor (86). Moreover, the phosphorylation sites necessary for arrestin binding can be localized in the receptor C-terminus (e.g., rhodopsin (83), NTSR1 (87)), ICL3 (e.g., M1 (88) and M2 muscarinic receptor (56, 89), 5-HTR1B (87)), or even ICL2 (e.g., μ-opioid receptor (90)). In fact, this complex and variable phosphorylation-dependent regulation led to the “barcode hypothesis”, the idea that different phosphorylation patterns on receptors can direct arrestin-dependent signaling (51-53). In the context of recent structural data, one mechanism consistent with barcode hypothesis is that different receptor phosphorylation patterns induce distinct conformations of bound arrestin, which in turn promote binding of different effectors (62).

We evaluated how the phosphorylation pattern in the D1R relates to directing signaling toward ERK1/2 or Src. We found that arrestin binding to peptides representing the C-terminus faithfully recapitulates existing data (compare (38, 55) and **Fig. 2**). Although the C-terminus contributes to arrestin recruitment to D1R, arrestin-3 can bind Class A GPCRs lacking C-terminal phosphorylation sites (59). We also observed some binding in a triple alanine D1R mutant, D1R^T428A/S431A/T439A^ (**Fig. 4**)

We found that phosphorylation of ICL3 regulates signaling **(Fig. 3)**. Prior work by Kim et. al. showed that ICL3 is the main regulatory region for D1R-arrestin interaction and receptor desensitization (38). Our results add new dimension, showing that D1R-ICL3 is not only important for arrestin interaction but regulates the direction of signaling. Specifically, we found that the phosphorylation of S258 and S259 of ICL3 promotes signaling toward ERK1/2, but not to Src **(Fig. 5C and D, 6B and F)**.

In conclusion, our results (summarized in **Fig. 8)** support the main premise of the barcode hypothesis: receptor phosphorylation pattern dictates the direction of arrestin-mediated signaling. Importantly, these results extend the barcode hypothesis and show that the phospho-barcode also impacts G protein coupling. In this study, we focused on the D1R-ICL3 region and did not screen all of the possible phosphorylation combinations in the D1R-ICL3, D1R-C-terminus or all branches of arrestin-mediated signaling. Therefore, we cannot rule out that D1R mediated signaling is influenced by the C-terminus, either alone or in combination with D1R-ICL3. Nevertheless, it is tempting to hypothesize that other residue combinations bias arrestin-mediated signaling in this system towards different effectors (7, 8). This is consistent with reported different shapes of the arrestin-receptor complex (68-70). We also identified different effects of G protein on this process, with Gs enhancing ERK1/2 but not Src activation. These findings help to better identify the origins of the signaling code by GPCRs.

**Figure 8.**
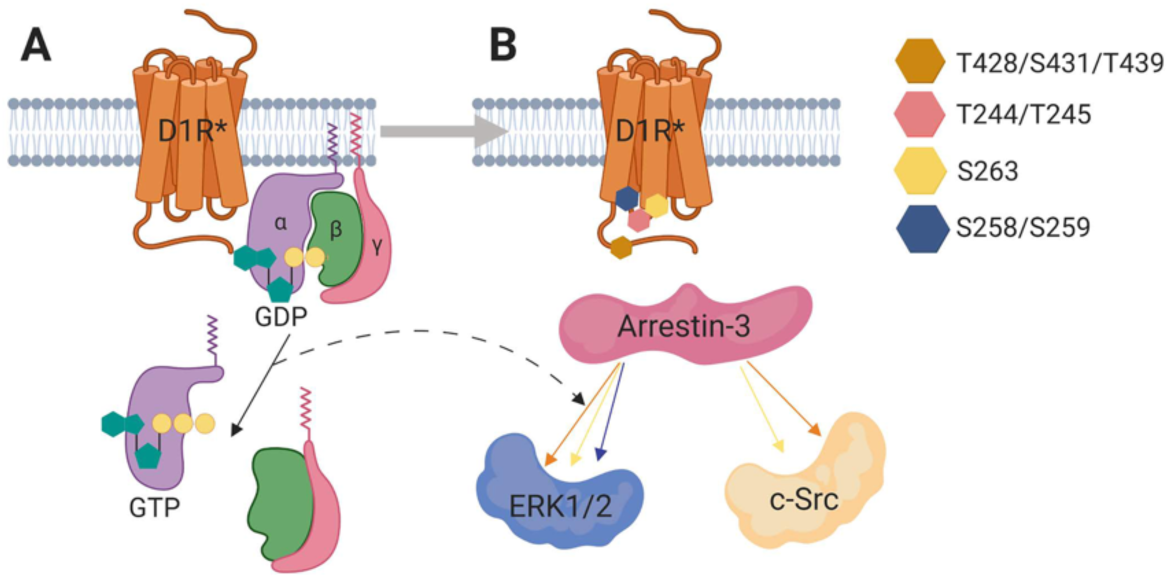
Model of biased signaling from the D1R receptor. Our data suggests that D1R-mediated ERK1/2 and Src activation depends on arrestin, and that Gαs is not obligatory. However, Gαs enhances ERK1/2 activation in a D1R-dependent fashion. The mechanism is likely a phospho-barcode-dependent development of a population of D1R-arrestin-3 and D1R-Gαs complexes in the cell, with activated Gαs enhancing ERK1/2 signaling downstream of active arrestin. Phosphomimetics in ICL3 affect Gαs activation, while phosphomimetics in the C-terminus do not. We further categorized phosphorylated residues of ICL3 into three groups: 1) T244 and T245 of ICL3 do not contribute to D1R mediated ERK1/2 or Src activation; 2) S258 and S259 residues significantly affect ERK1/2 but not Src phosphorylation, which indicates that phosphorylation of these residues stabilizes an arrestin-3 conformation that contributes to ERK1/2 signaling; 3) S263 affects both ERK1/2 and Src activation. Figure made using BioRender (BioRender.Com).

## Materials and Methods

Detailed procedures for the protein expression and purification, and all biochemical studies are described in SI Appendix, SI Materials and Methods. All data discussed in this study are included in the manuscript or SI Appendix.

## Acknowledgments

We are grateful to Dr. Asuka Inoue for the arrestin-2/3 and Gs KO HEK293 cells and Dr. Jonathan A. Javitch for pcDNA3.1-SF-hPTX plasmid.

## Author Contributions

AIK, TMI and VVG designed research; AIK and NAP performed experiments; AIK analyzed data; AIK, NAP, TMI and VVG wrote the paper. All authors approved the final version of the manuscript.

## Figures

**Supplementary Figure 1:**
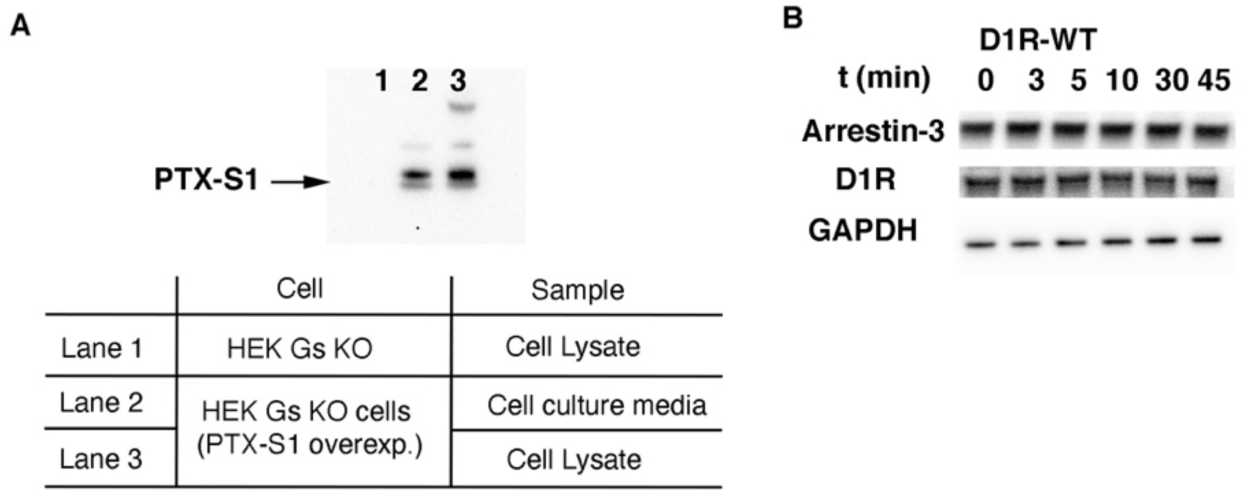
Pertussis toxin, Arrestin-3, D1R and GAPDH expression in Gs/Golf KO HEK cells. (A) Gs/Golf KO HEK-293 cells co-expressed with D1R, arrestin-3 and PTX. We used pcDNA3.1-SF-hPTX plasmid to express signal peptide and FLAG-tagged PTX S1 subunit in Gs KO HEK-293. After expression, PTX protein is secreted outside of the cell and then taken up from the cell culture media as described (14). PTX expression was detected with an anti-FLAG antibody and Western blotting. (Lane 1) The PTX expression in Gs/Golf KO HEK-293 cell lysate. Lane (2) and (3) are showing Gs/Golf KO HEK-293 cells expressed with PTX. Lane 2 is the sample from cell culture media, Lane 3 is the cell lysate. (B) We used a pan arrestin antibody to detect arrestin-3, an HA antibody to detect D1R, and a GAPDH antibody as an internal control for each sample in our phosphorylation assays. Data repeated with at least three independent experiments and representative images are shown.

**Supplementary Figure 2.**
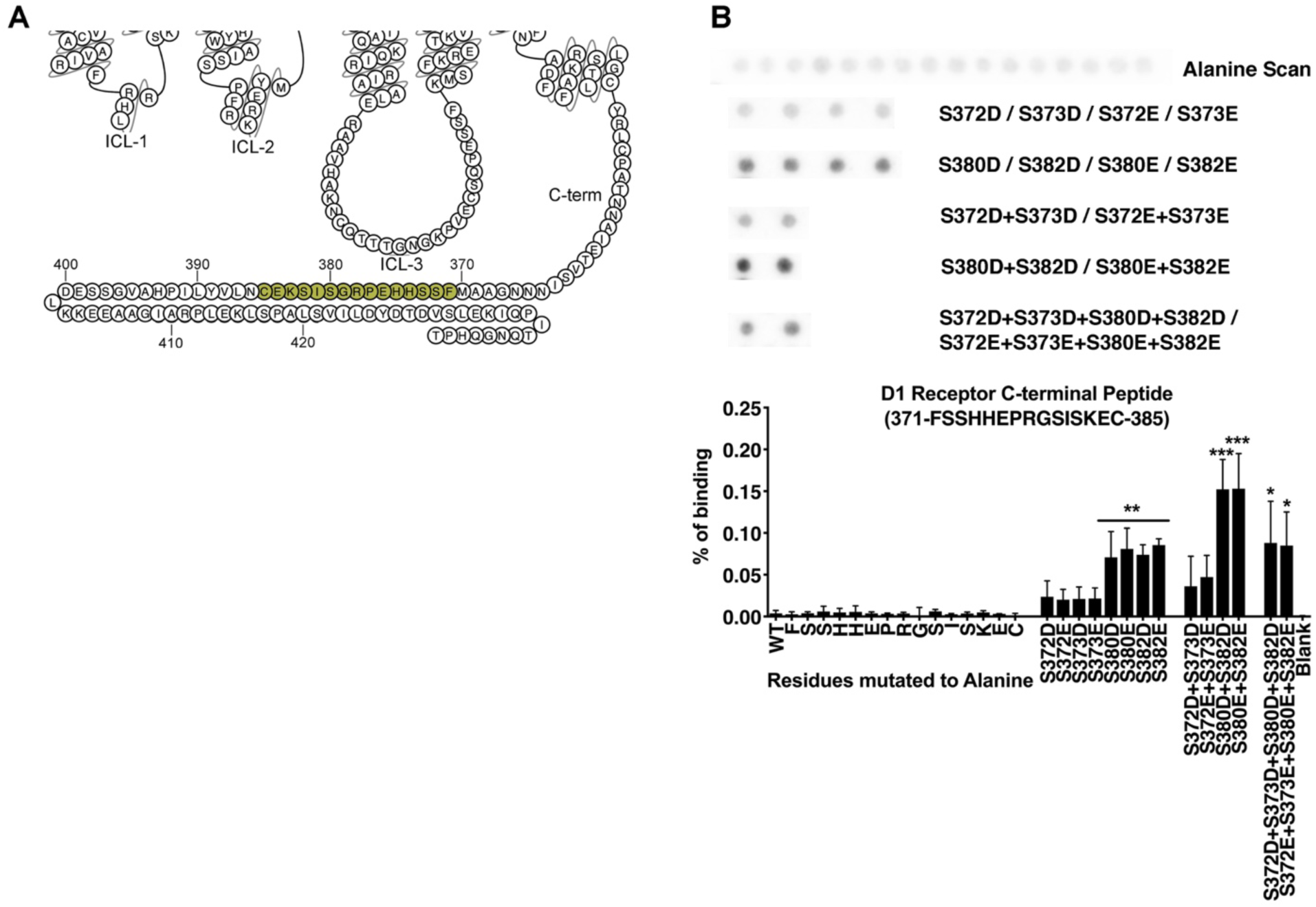
Peptide array analysis for the D1R F371-C385 peptide. (A) A snake diagram (1) highlighting the residues that were tested for their role in arrestin-3 binding. (B) Peptide array analysis of the D1R^371-385^ peptide. As monitored by Far Western analysis (see Methods), the wild-type peptide exhibited little detectable interaction with arrestin-3; alanine scanning did not significantly impact arrestin-3 binding. Phosphomimetic substitution at positions S380 and S382 increase binding, with an additive effect. However, phosphomimetic substitution of S372 and S373 interfered with this interaction. Data represent at least three independent experiments. Representative peptide array results are shown. Statistical analysis was performed using one-way ANOVA followed by Tukey’s post-hoc test. The statistical significance of the difference of the signal from the WT peptide spot is shown, as follows: *, p<0.05; **, p<0.01; ***, p<0.001.

**Supplementary Figure 3:**
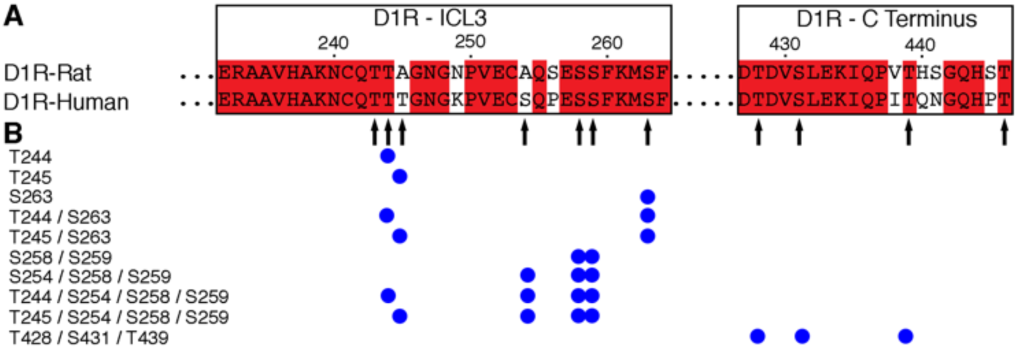
Mutagenesis of the D1R intracellular loop 3 and C-terminus. (A) Sequence alignment of the rat and human D1R in the ICL3 and C-terminus. Arrows indicate S/T residues in the ICL3 and the C-terminus of human D1R. (B) Summary of mutations investigated. Blue circles indicate the locations of mutations to alanine or phosphomimetics.

**Supplementary Figure 4:**
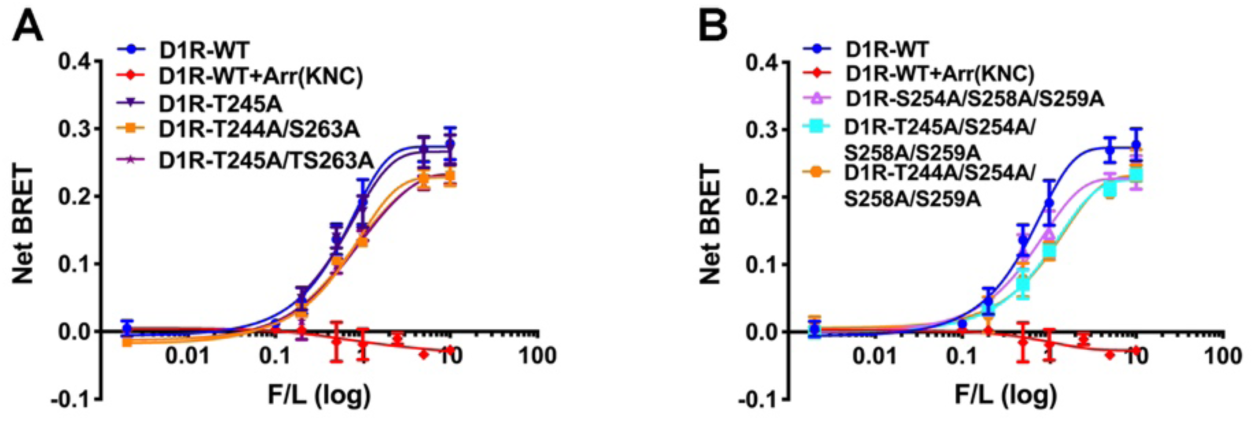
In-cell binding of arrestin-3 to WT and mutant D1R. BRET assays were used to measure the binding of wild type and mutant Rluc8-D1Rs to Venus-Arrestin-3 in arrestin2/3 knock-out HEK293 cells (A-B). Mutant D1R receptors were screened in the presence of increasing amounts of Venus-arrestin3 (0-1µg) along with 50–100 ng of the plasmid encoding indicated D1R. KNC arrestin (Arr(KNC)) was used as a negative control as it is not capable of binding to receptors (1). The net BRET ratio was calculated as the acceptor emission divided by the donor emission and expressed as the relative change compared with unstimulated cells as describe in the method section. Data represent 3-5 independent experiments.

**Supplementary Figure 5:**
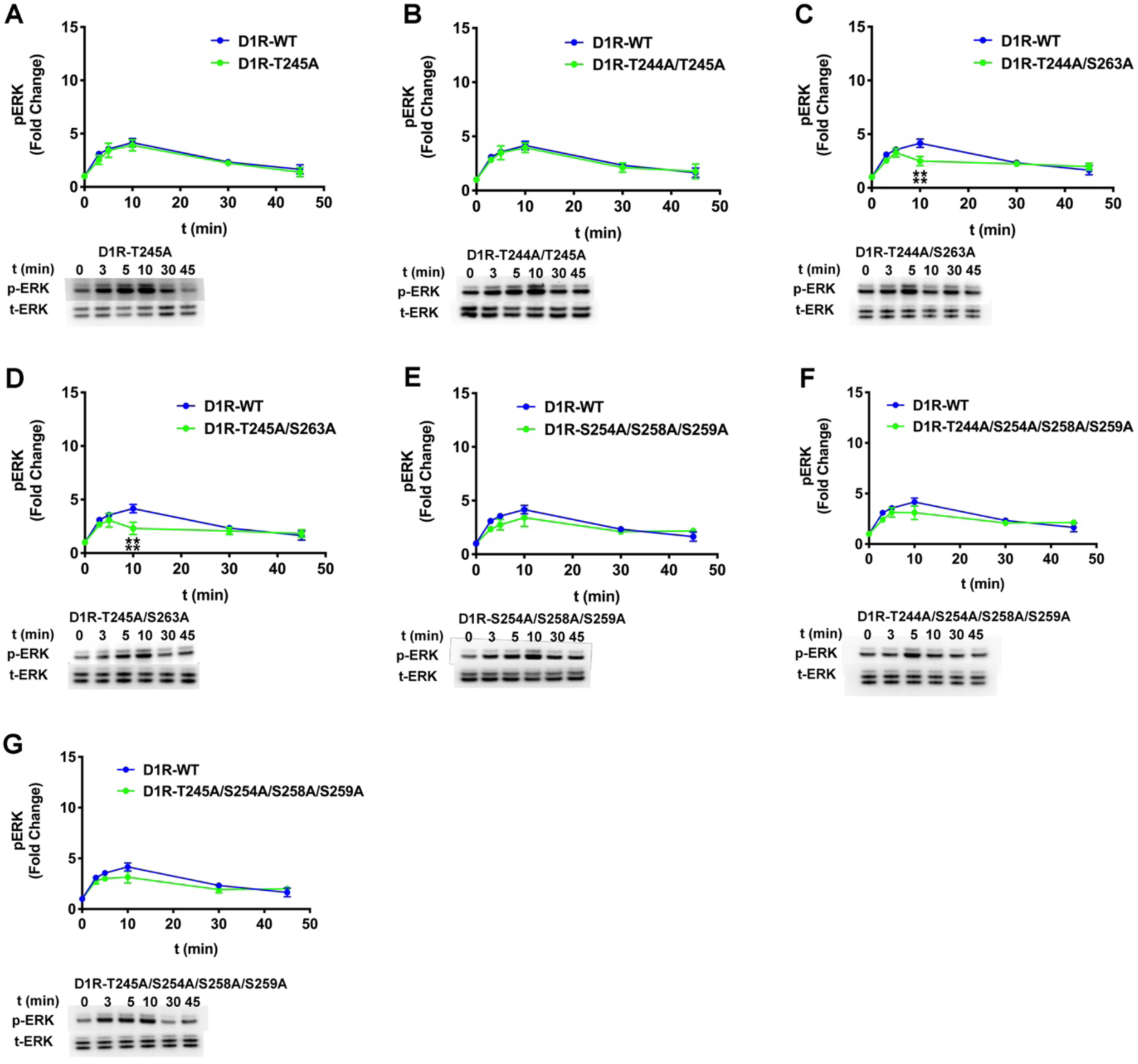
D1R phosphorylation pattern affects activation of ERK1/2. Arr2/3 KO HEK-293 cells were overexpressed with D1R and Arrestin-3. The assay was initiated by adding 10 µM Dopamine at 37 °C for the indicated time points. Both phosphorylated ERK (pERK) and total ERK (tERK) were visualized by Western blotting. Cell lysates were separated on 4-12% gradient SDS-PAGE, transferred to PVDF membranes and immunoblotted using the indicated antibody. Signals were quantified by densitometry and expressed as the fold change of unstimulated samples in each experiment. Representative images are shown for each receptor mutant group. Data represents at least three independent experiments. Statistical analysis was determined with one-way ANOVA followed by Tukey’s post hoc test with correction for multiple comparisons between WT- and mutant receptor mediated signal in each time points (***P<0.001).

**Supplementary Figure 6:**
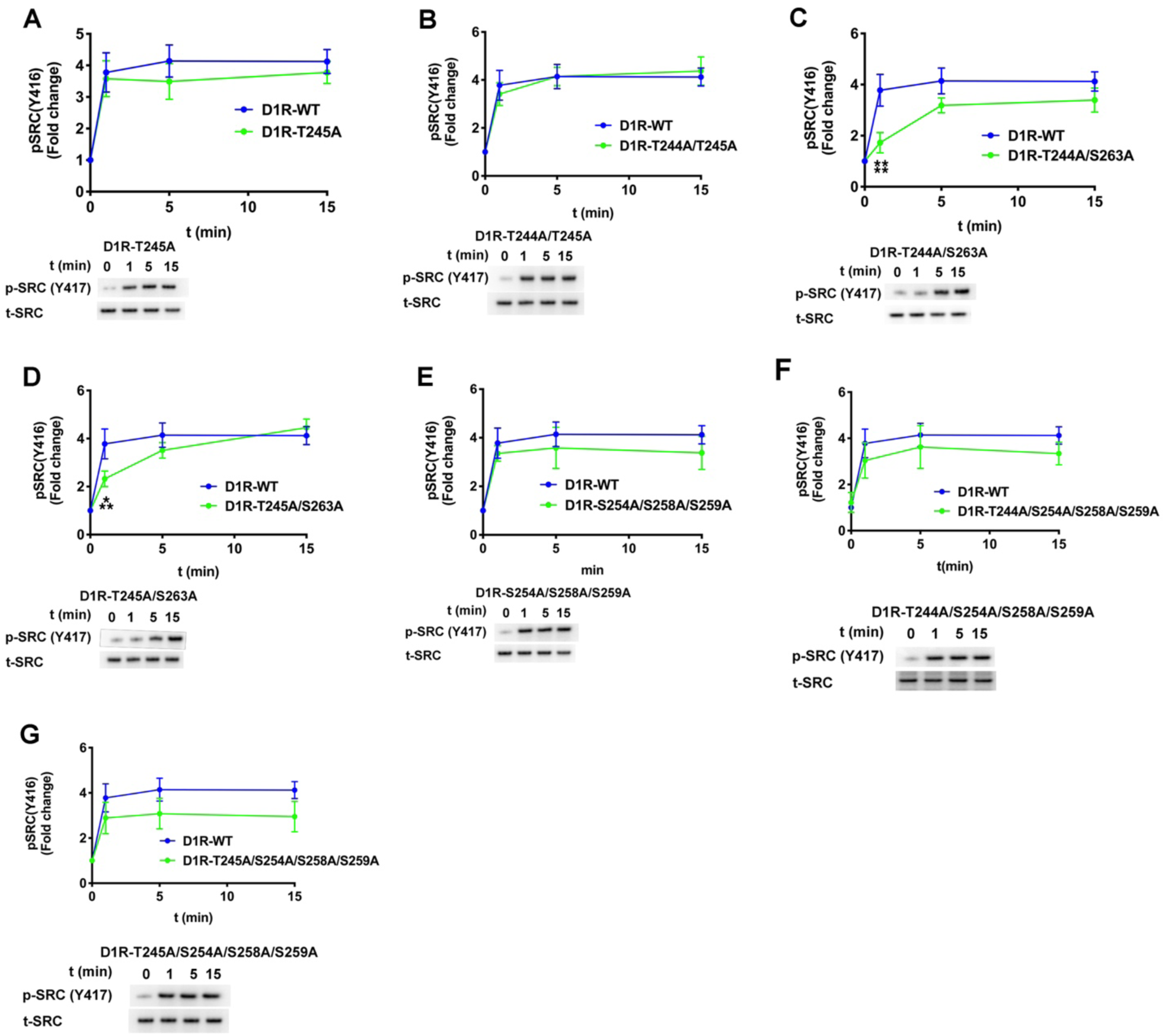
The effect of phosphorylation pattern of D1R on the Src activation. Arr2/3 KO HEK-293 cells were overexpressed with D1R and arrestin-3. The assay was initiated by adding 10 µM Dopamine at 37 °C for indicated time points. Both phosphorylated SRC (pSRC) and total SRC (tSRC) were visualized by Western blotting. Cell lysates were separated on 4-12% gradient SDS-PAGE, transferred to PVDF membranes and immunoblotted using the indicated antibody. Signals were quantified by densitometry and expressed as the fold change of unstimulated samples in each experiment. Representative images are showing from each receptor mutant group. Data represents at least three independent experiments. Statistical analysis was determined with one-way ANOVA followed by Tukey’s post hoc test with correction for multiple comparisons between WT- and mutant receptor mediated signal in each time points (***P<0.001; ****P<0.0001).

**Supplementary Figure 7:**
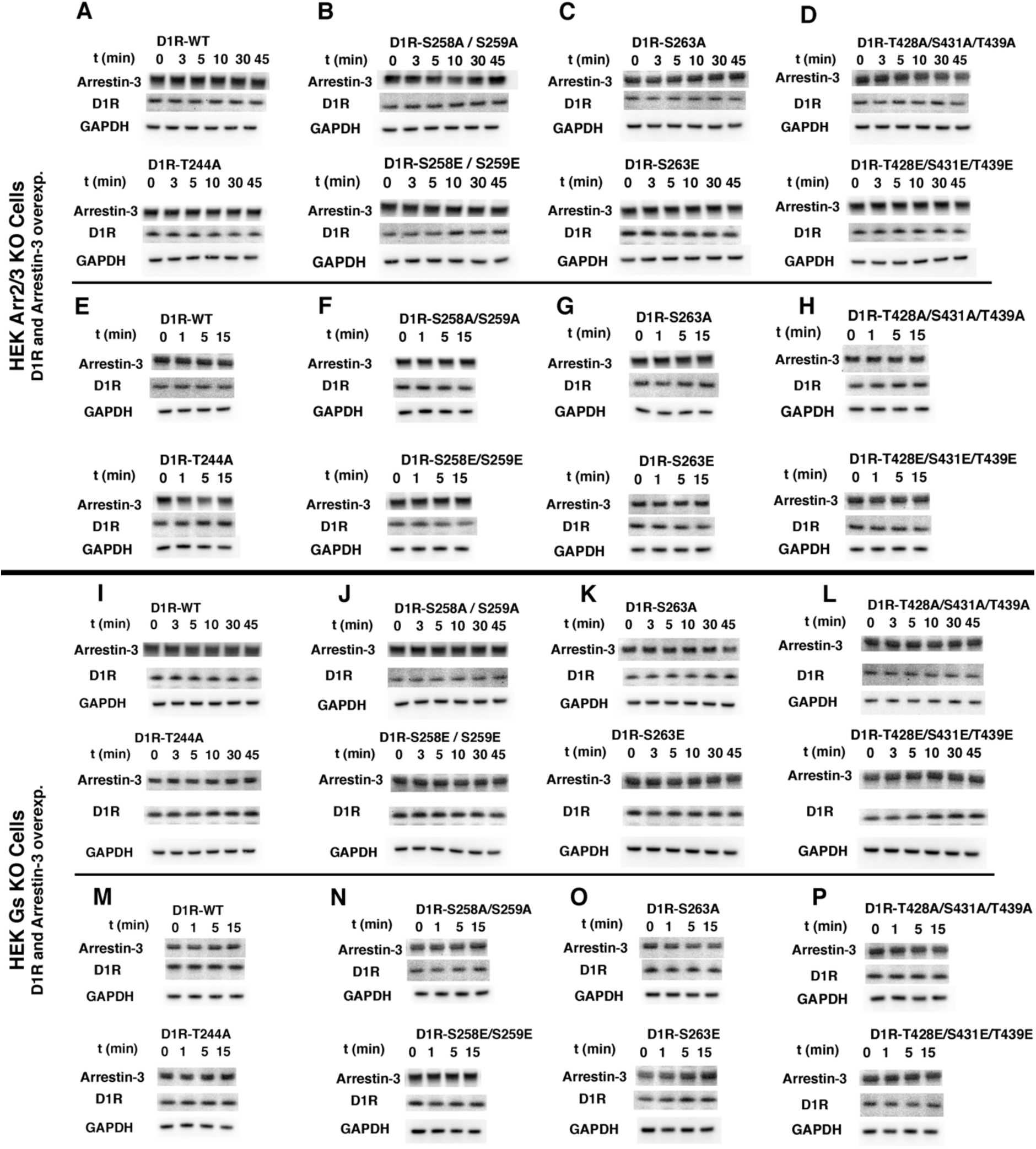
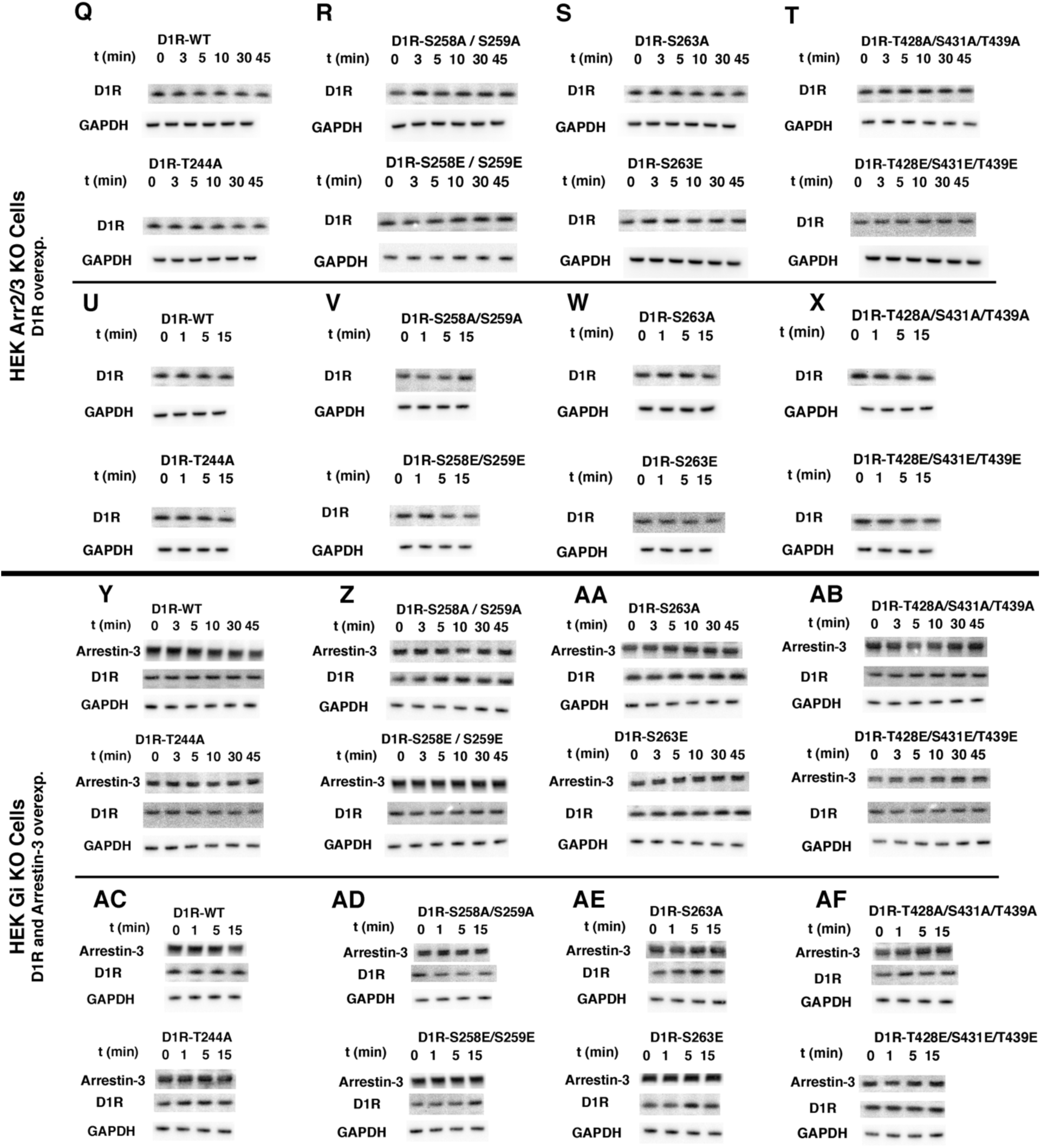
Arrestin-3, D1R and GAPDH expression levels in HEK cells. We used an arrestin antibody to detect arrestin-3, HA antibody to detect D1R and GAPDH antibody as an internal control for each sample in our phosphorylation assays. The samples from the Arr2/3 KO HEK-293 cells co-expressed with D1R and Arrestin-3, the Gs KO HEK-293 cells co-expressed with D1R and the Arrestin-3, Arr2/3 KO HEK-293 cells expressed with D1R, Gi KO HEK-293 cells co-expressed with D1R and the Arrestin-3 are showing in the first, second, third and fourth panel, respectively. GAPDH detected with anti-GAPDH antibody by using Western blotting. Data repeated with at least three independent experiments and representative images are showing from each sample.

**Supplementary Figure 8:**
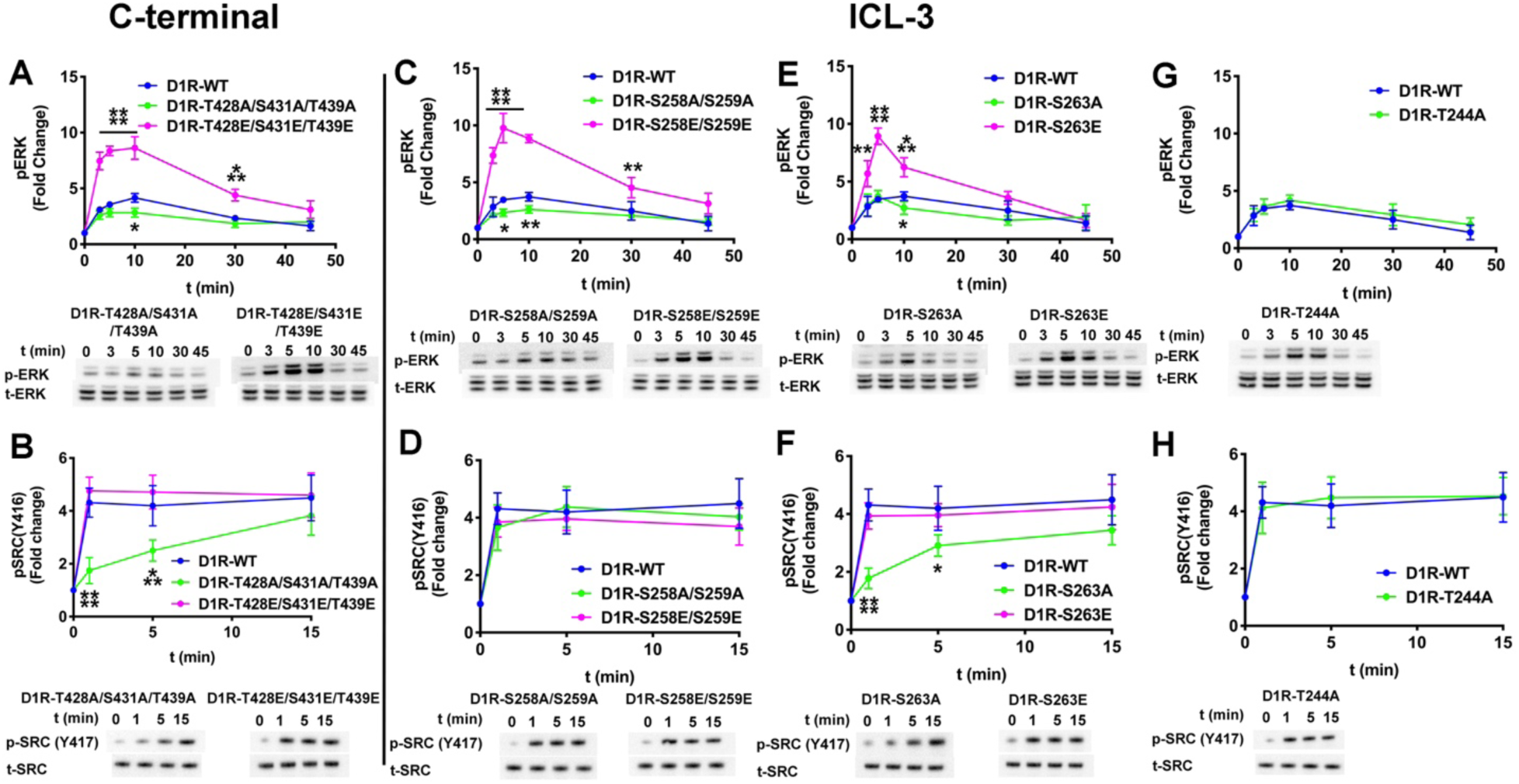
The effect of the D1R-ICL3 phosphorylation pattern on ERK1/2 and Src kinase phosphorylation. Gαi/o/t KO HEK-293 cells were co-transfected with plasmids encoding arrestin-3 and either wild-type or variant D1R. The assay was initiated by adding 10 µM dopamine at 37 °C. Cell lysates were separated on a 4-12% gradient SDS-PAGE, transferred to PVDF membranes and immunoblotted for phosphorylated ERK1/2 (pERK), phosphorylated Src (pSrc), total ERK1/2 (tERK), and total Src (tSRC). Signals were quantified by densitometry and expressed as the fold change relative to unstimulated samples in each experiment. (A) – (D) Time course of ERK 1/2 phosphorylation associated with dopamine stimulation of cells expressing wild-type or variant D1R. (E) – (H) Time course of Src kinase activation associated with dopamine stimulation of cells expressing wild-type or variant D1R. Data represent at least three independent experiments. GAPDH expression was used as an internal control for each sample (**SI Appendix Fig. S6**). Statistical analysis was performed with one-way ANOVA followed by Tukey’s post hoc test with correction for multiple comparisons between wild-type and mutant D1R mediated signal at each time point. Significance is shown, as follows: *, p<0.05; **, p<0.01; ***, p<0.001; ****, p<0.0001.

**Supplementary Figure 9:**
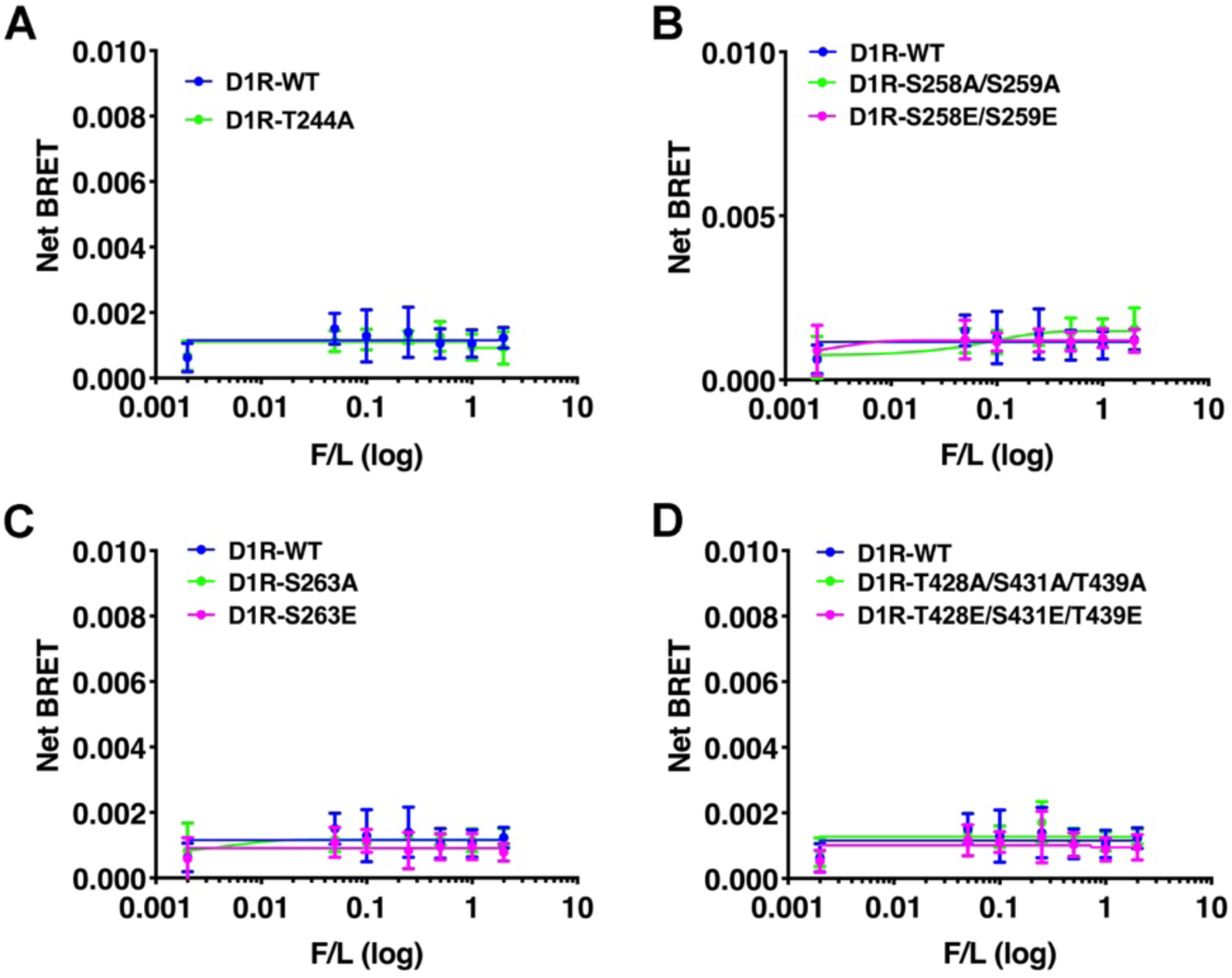
In-cell proximity between arrestin-3 and Gαs suggests against formation of a D1R-Gs-arrestin-3 super complex. A BRET assay was used to measure the proximity between Rluc8-Arrestin3 and Gαs-YFP in the presence of WT and mutant D1R in the arrestin-2/3 KO HEK293 cells as described in method section. (A) – (D) BRET signal of wild-type and variant D1R receptors stimulated by 10 µM dopamine in the presence of increasing amounts of Gαs-YFP (0-1µg) and monitored for 10 min. The net BRET ratio was calculated as the acceptor emission divided by the donor emission and expressed as the relative change compared with unstimulated cells. Statistical analysis was performed using one-way ANOVA followed by Tukey’s post-hoc test and we didn’t find any statistical differences.

**Supplementary Figure 10:**
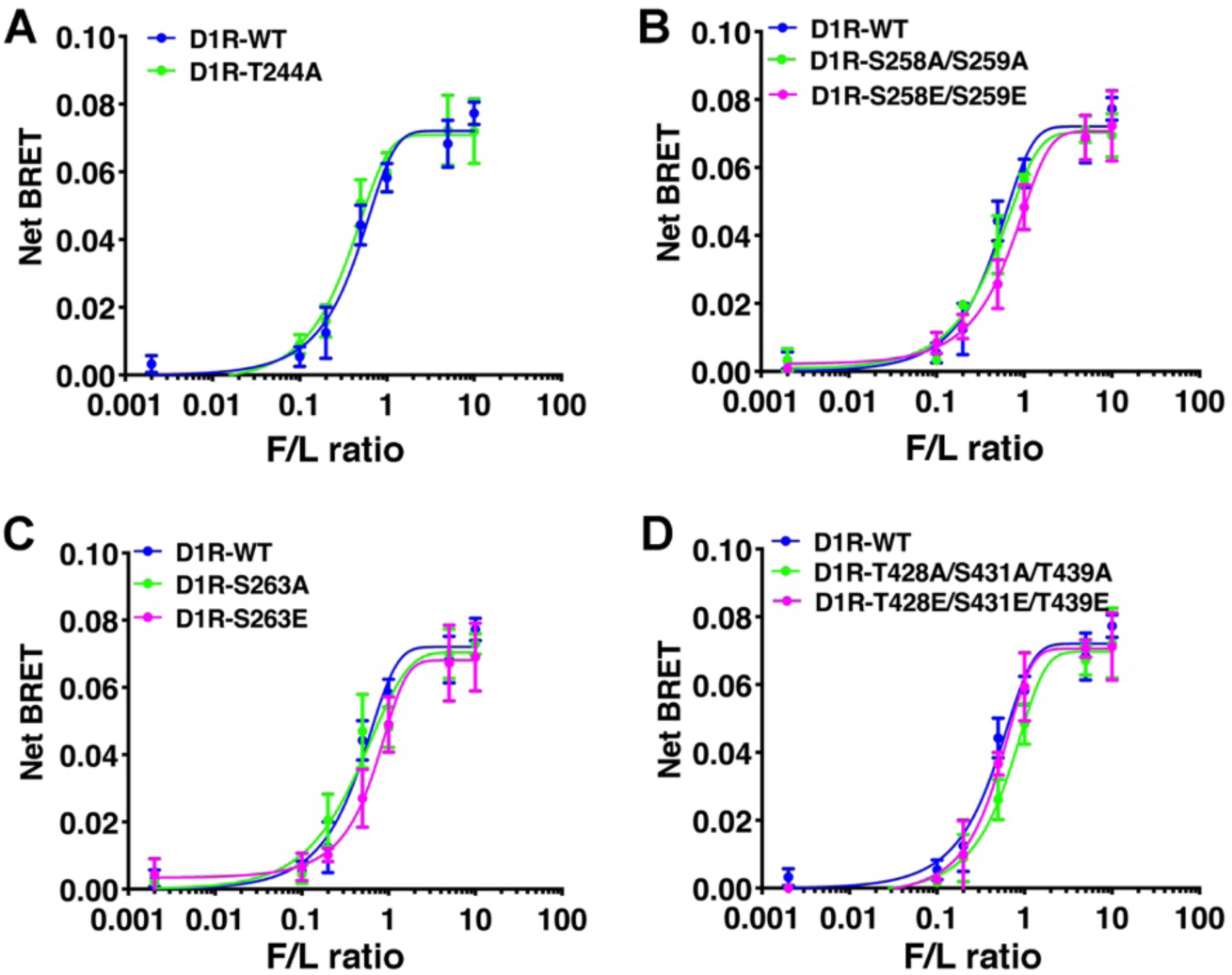
In-cell binding of Gαs to WT and mutant D1R in the absence of arrestin. A BRET assay was used to measure the binding between WT and mutant Rluc8-D1R to Gαs-YFP in arrestin-2/3 KO HEK293 cells as described in the legend to Fig. 4. (A) – (D) BRET signal of wild-type and variant D1R receptors stimulated by 10 µM dopamine in the presence of increasing amounts of Gαs-YFP (0-1µg) and monitored for 10 min. The net BRET ratio was calculated as the acceptor emission divided by the donor emission and expressed as the relative change compared with unstimulated cells. Statistical analysis was performed using one-way ANOVA followed by Tukey’s post-hoc test and we didn’t find any statistical differences.

**Supplementary Figure 11:**
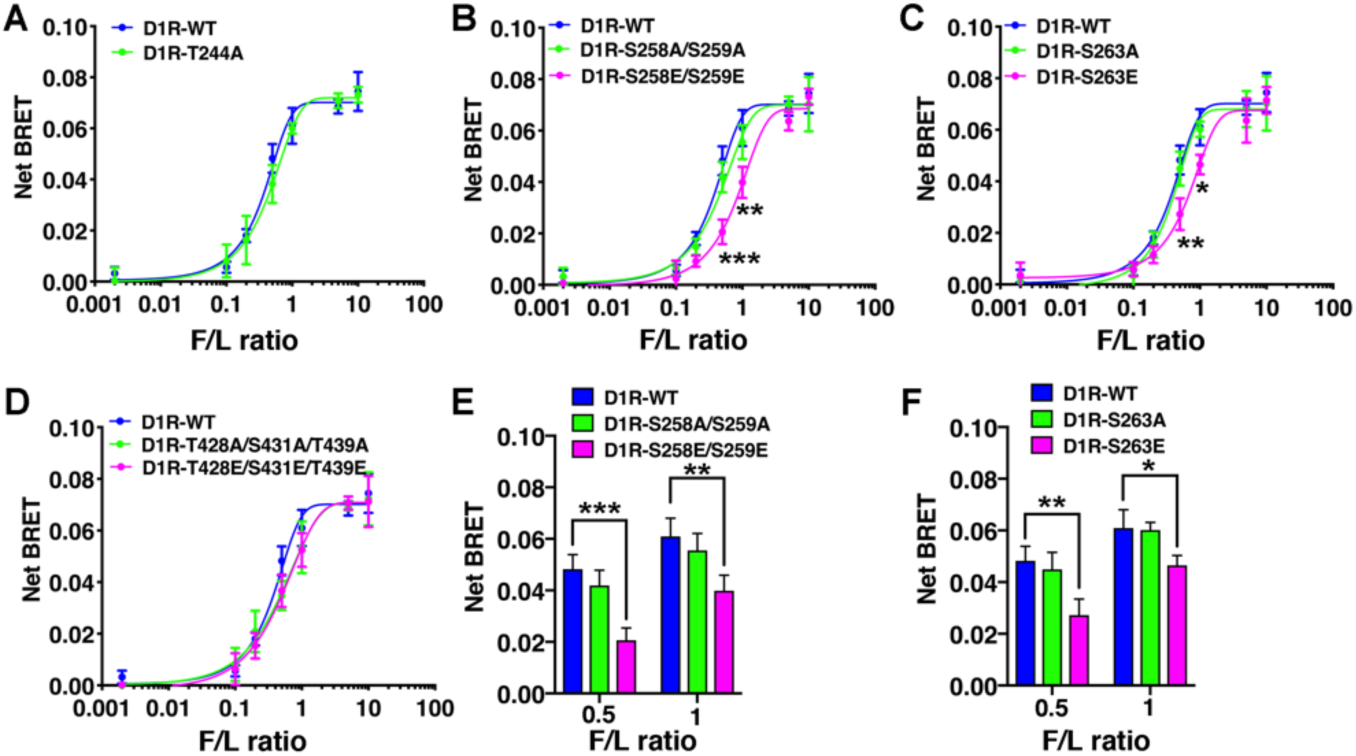
In-cell binding of Gαs to WT and mutant D1R in the presence of overexpressed arrestin-3. A BRET assay was used to measure the binding between WT and mutant D1R-Rluc8 to Gαs-YFP. (A) – (D) BRET signal of wild-type and mutant D1R stimulated by 10 µM dopamine in the presence of increasing amounts of Gαs-YFP (0-1µg) monitored for 10 min. The net BRET ratio was calculated as the acceptor emission divided by the donor emission and expressed as the change relative to unstimulated cells. Statistical analysis was performed using one-way ANOVA followed by Tukey’s post-hoc test. (E) The statistical significance of the signal detected at the indicated donor acceptor ratios between the WT-D1R and the D1R-S258E-S259E (**, p<0.01, ***, p<0.001). (F) The statistical significance of the signal detected at the indicated donor acceptor ratios between the WT-D1R and the D1R-S263E (*, p<0.05, **, p<0.01).

## Footnote

1 We use systematic names of arrestin proteins, where the number after the dash indicates the order of cloning: arrestin-1 (historic names S-antigen, 48 kDa protein, visual or rod arrestin), arrestin-2 (β-arrestin or β-arrestin1), arrestin-3 (β-arrestin2 or hTHY-ARRX), and arrestin-4 (cone or X-arrestin).

