## Supplementary material for "Phosphorylation barcode-dependent signal bias of the dopamine D1 receptor": SI Materials and Methods

### Supplementary Information

#### Materials and Methods

*Materials.* The Superdex S200 Increase 10/300 GL column was from GE life sciences (Marlborough, MA). Primary antibodies Total-ERK1/2 (9102S), pERK1/2 (9101S), Src (2109S), pSrc-T416 (6943S), anti-Flag (2368) were from Cell Signaling Technology (Danvers, MA), F431 antibody used to detect Arrestin protein (1), pan-Gi (SC-26761) was from Santa Cruz Biotech., Gs (n192/12) was from EMP Millipore Burlington, MA), HA-tag antibody was from (16B12) Covance (Dedham, MA), GAPDH (AB2302) was from Millipore-sigma (Burlington, MA). HRP-conjugated secondary antibodies (anti-rabbit: 5520-0337, anti-mouse: 55220-0341) were from Seracare (Milford, MA) and anti-goat (sc-2354) was from Santa Cruz Biotech (Dallas, TX). Cell culture media and fetal bovine serum were from Gibco (Waltham, MA). Dopamine hydrochloride was from Abcam (ab120565, Cambridge, MA). Enhanced chemiluminescence substrate (34580) was from ThermoFisher (Waltham, MA). Lipofectamine 2000™ was from Invitrogen (Waltham, MA).

*Arrestin-3-(1-393) expression and purification.* Arrestin-3 (1-393) was expressed and purified as described (1-5) followed by an additional purification step on a Superdex S200 Increase 10/300 GL column (GE Life Sciences, Pittsburgh, PA) equilibrated in 20 mM MOPS, pH 7.5, 150 mM NaCl, and 2 mM TCEP. Protein purity was assessed by SDS-PAGE with Coomassie staining. Arrestin-3 (1-393) concentrations were determined spectroscopically using corresponding molecular weight (44 kDa) and molar extinction coefficient ($16.5 \times 10^3$ $m^{-1}$ $cm^{-1}$) (Synergy Neo2 HTS Multi-Mode Microplate Reader, BioTek).

*Expression constructs.* For the BRET assays, pcDNA3 encoding Renilla luciferase variant 8 (RLuc8) fused to the D1 receptor C-terminus (6) served as the template for introducing mutations. For a acceptor partner of the BRET, Venus-arrestin fusion construct was used (6). For the ERK1/2 and Src activation assays, pcDNA3 encoding D1R-HA tag was used for introducing mutations. All mutations were confirmed by DNA sequencing (GenHunter DNA Sequencing Service, Nashville, TN).

*Peptide Array analysis and Far-western blot.* Peptide array synthesis was performed using the ResPep SL peptide synthesizer (Intavis AG, Koeln, Germany) according to standard SPOT synthesis protocols (7, 8). D1R-derived peptides containing 15 amino acids were directly coupled to membranes via the C-terminus during synthesis. Dried membranes with peptides were soaked in 100% ethanol for 5 min then rehydrated in water (twice 5 min). The membranes were blocked for 1 h in Tris-buffered saline (TBS) with 5% milk and 0.05% Tween 20 (Sigma-Aldrich, St. Louis, MO) and washed 3 times (5 min each) in TBS with 0.05% Tween 20 (TBS-T). The membranes were incubated overnight at 4°C with arrestin-3 (1-393) at a final concentration of 0.5 µM in 20 mM MOPS, pH 7.5 buffer containing 150 mM NaCl and 2 mM TCEP. The next morning, membranes were washed 3 times (5 min each) in TBS-T buffer and incubated with primary antibody against arrestin (F431, (9) at 1:5,000 dilution in TBS-T for 1 h). Spots were detected using HRP-conjugated secondary antibodies as described by the manufacturer. For densitometric analysis, the signal in individual dots was quantified as a percentage of total density detected on the membrane (Quantity One and Image Lab, Bio Rad (Hercules, CA), Image J (10)). This allowed for comparison of intensities across all peptide experiments. Statistical analysis was performed using one-way ANOVA followed by Dunnett's post-hoc test using GraphPad Prism 8.0.

*Cell Culture.* Arrestin 2/3, Gs and Gi KO HEK293 cells were a gift from Dr. Asuka Inoue (Tohoku University). WT and KO HEK293 cells were maintained in Dulbecco's modified Eagle's media (DMEM) (Invitrogen) supplemented with penicillin (100 IU/ml), streptomycin (100 µg/ml), and 10% (v/v) fetal bovine serum, at 37 °C and 5% $CO_2$. HEK-293 cells were transfected using a 1:2 DNA:Lipofectamine 2000 (Invitrogen) ratio according to the manufacturer's instructions.

*BRET assay.* BRET was performed as described previously (6, 11). In brief, the donor was Renilla luciferase variant 8 (RLuc8) C-terminally fused to wild-type or mutant D1R in pcDNA3.1 (6). The acceptor was arrestin-3 with N-terminal Venus in pcDNA3.1(6). HEK-293 cells (80-90% confluent) were co-transfected with Venus-arrestin (0-1µg), wild-type or mutant RLuc8-D1R (50 ng), and empty pcDNA3.1 (to equalize DNA). The agonist (10 µM dopamine) was added prior to the addition of 5 µM coelenterazine-*h* (12). The net BRET ratio was calculated as the long wavelength emission (530 nm) divided by the short wavelength emission (480 nm) and expressed as the relative change compared with unstimulated cells. The Venus-arrestin-3 fluorescence, which is directly proportional to the expression level, was normalized by the basal luminescence from the respective D1R-RLuc8 construct (F/L ratio) to account for variations in cell number and expression. For the D1R and Gαs interaction, Rluc8-D1R was used as donor and Gαs-YFP was used as an acceptor. Gαs-YFP was a gift from Dr. Catherine Berlot (Addgene plasmid #55781) (13). For the Gαs and arrestin3 interaction, Rluc8-arrestin3 was used as donor and Gαs-YFP was used as an acceptor. Net BRET data were fitted to a one-site binding hyperbola or a simple linear regression (when convergence was not achieved). Curve fits and statistical significance ($p < 0.05$) were determined using a Student's t-test or One-Way Analysis of Variance with Tukey's multiple comparison test where appropriate using GraphPad Prism 8.0.

*Phosphorylation Assay and Immunoblotting.* For the ERK1/2 and Src activation assays, wild-type and mutant D1R with a N-terminal HA tag in pcDNA3.1 was co-transfected with arrestin-3 with a C-terminal HA tag in arrestin-2/3 KO or Gs/G-olf KO HEK293 cell lines. All experiments were conducted 48 h after transfection. For the pertussis toxin expression, Gs/Golf KO HEK293 cells were co-transfected with D1R, arrestin-3, and pcDNA3.1-sf-PTX plasmid 24 h before the experiment (14). In this construct PTX S1 subunit is fused with both a signal peptide and a FLAG tag. After expression, the pertussis toxin is secreted from the cytoplasm and taken back up from the cell media. Cells were serum starved for 16 h and the assay was initiated by adding 10 μM dopamine. The medium was removed and 400 μl 2x Laemmli buffer was added. Samples of equal volume from the cell lysates were separated on 4-12% gradient SDS-PAGE, transferred to PVDF membranes and immunoblotted. Signals were quantified by densitometry and expressed as the fold change of unstimulated samples in each experiment. Western Blot images were taken using Biorad ChemiDoc system (BioRad Life Sciences, Hercules,CA). Files were cropped and converted to a JPEG or TIFF file using BioRad Image lab software (BioRad Life Sciences, Hercules,CA). Each figure panel was aligned using Adobe Photoshop CS6 software. Statistical analysis was performed using One-way ANOVA followed by Tukey's post-hoc test for multiple comparisons.
